# Positive and biphasic extracellular waveforms correspond to return currents and axonal spikes

**DOI:** 10.1101/2023.03.24.534099

**Authors:** Shirly Someck, Amir Levi, Hadas E. Sloin, Lidor Spivak, Roni Gattegno, Eran Stark

## Abstract

Multiple biophysical mechanisms may generate non-negative extracellular waveforms during action potentials, but the origin and prevalence of positive spikes and biphasic spikes in the intact brain are unknown. Using extracellular recordings from densely-connected cortical networks in freely-moving mice, we find that a tenth of the waveforms are non-negative. Positive phases of non-negative spikes occur in synchrony or just before wider same-unit negative spikes. Narrow positive spikes occur in isolation in the white matter. Isolated biphasic spikes are narrower than negative spikes, occurring right after spikes of verified inhibitory units. In CA1, units with dominant non-negative spikes exhibit place fields, phase precession, and phase-locking to ripples. Thus, near-somatic narrow positive extracellular potentials correspond to return currents, and isolated non-negative spikes correspond to axonal potentials. Identifying non-negative extracellular waveforms that correspond to non-somatic compartments during spikes can enhance the understanding of physiological and pathological neural mechanisms in intact animals.

## Introduction

Extracellular recordings have been employed extensively during the last century to monitor the electrical activity of multiple neurons^1, 2^. During an action potential, the recorded waveforms depend on neuronal morphology, spike initiation site, and the relative position of the electrodes and the subcellular compartments^3–7^. Extracellular potentials recorded next to different compartments are expected to exhibit distinct polarities^8–10^. However, most studies focus on somatic signals, recorded as negative deflections in an un-inverted extracellular record^11–14^, and non-negative potentials are rarely reported in vivo.

For simplified neuronal morphologies, computational models show that the somatic inward currents during a spike are balanced by outward “return currents” evident as positive extracellular spikes (P-spikes; **Fig. 1**), expected to be generated next to the proximal dendrites^5, 9, 15, 16^. P-spikes were reported to correspond to backpropagation of somatic spikes into the dendrites^17, 18^ and to outward currents in the dendrites^19–21^. Others reported that P-spikes are generated by axonal potentials^22, 23^. Near the axon, a propagating sodium spike is expected to yield inward currents with a negative phase, preceded by a wavefront of local return currents yielding a positive phase^9, 15^. Together, the two phases are expected to generate a biphasic spike (B-spike; **Fig. 2**). B-spikes were reported to correspond to axonal spikes^7, 22, 24–31^ or dendritic spikes^18, 32^. When sodium spikes initiate at the distal dendrites, P-spikes are expected next to the soma, and B-spikes at the proximal dendrites^21, 33^. Experimental studies suggested that B-spikes correspond to backpropagation of somatic spikes^34–36^.

**Figure 1.**
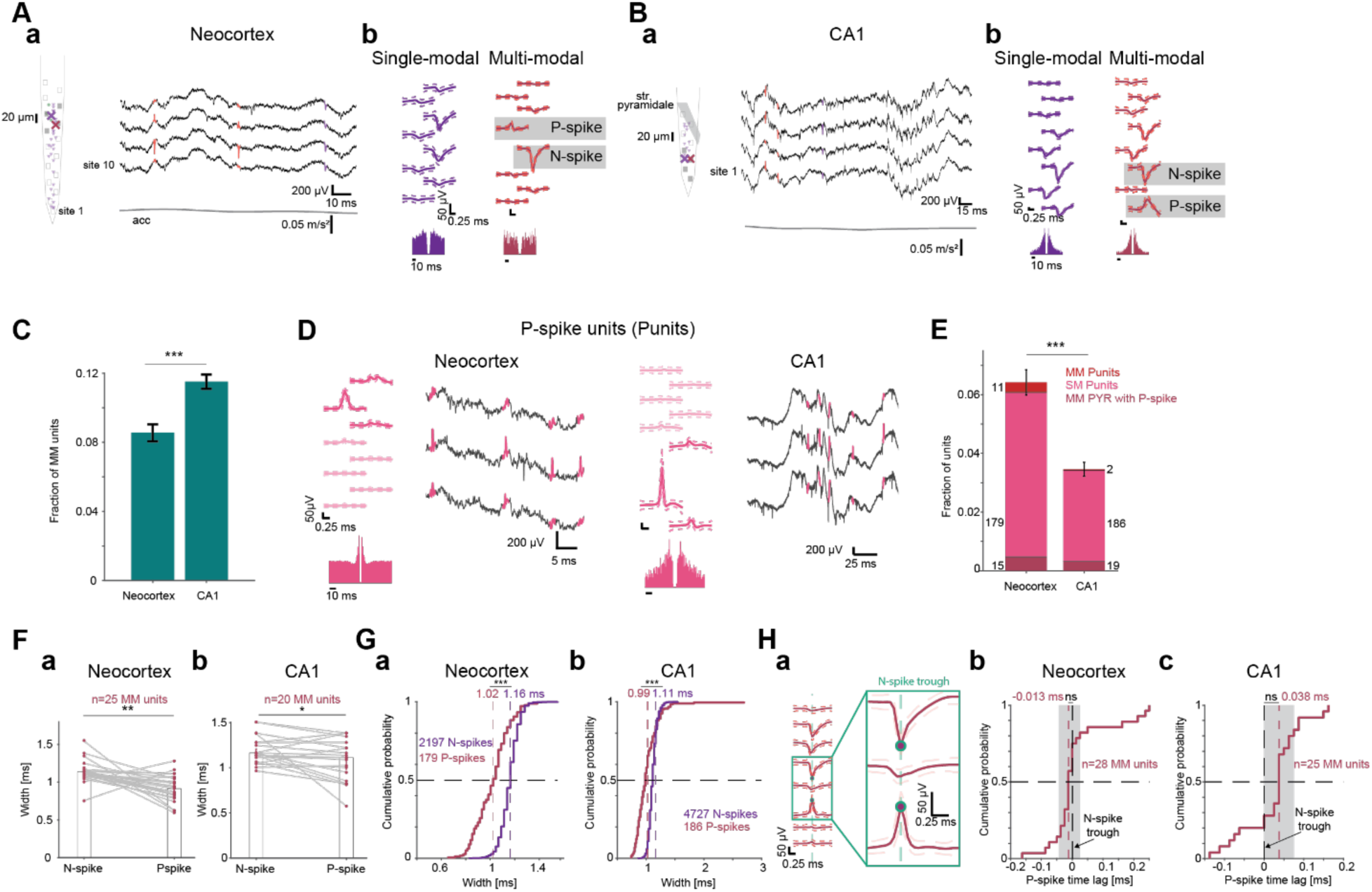
Positive extracellular spikes correspond to return currents. **(A)** Single-modal (SM) and multi-modal (MM) units recorded simultaneously from the neocortex of a freely-moving mouse. (**a**) **Left**, Schematic shank with 32 simultaneously-recorded units. X marks correspond to the units depicted in **b**. **Right**, Wideband (0.1-7,500 Hz) traces recorded by four adjacent electrodes. acc, head acceleration. (**b**) Wideband spike waveforms and autocorrelation histograms (ACHs) of the SM pyramidal cell (PYR) and the MM PYR with a negative spike (N-spike) and a positive spike (P-spike). **(B)** MM units appear in extracellular recordings from hippocampal region CA1. (**a**) **Left**, Shank schematic with 13 simultaneously-recorded units. **Right**, Wideband traces during a spontaneous ripple in CA1. (**b**) Example SM and MM units. **(C)** MM units are more prevalent in CA1 than in neocortex. Dataset includes 3193 neocortical units from n=17 mice and 5986 CA1 units from n=9 mice. Here and in **E**, ***: p<0.001, G-test. **(D)** Positive units (Punits) in extracellular recordings. A Punit is defined as a unit with a dominant positive peak in the main channel waveform. **(E)** Units with P-spikes are more prevalent in neocortex than in CA1. **(F)** P-spikes are narrower than same-unit N-spikes. Spike width, inverse of the dominant frequency of the waveform spectrum. MM PYRs with P-spikes and MM Punits are included. */**: p<0.05/p<0.01, Wilcoxon’s test. **(G)** P-spikes of SM Punits are narrower than N-spikes of SM PYRs or MM PYRs with B-spikes. ***: p<0.001, U-test. **(H)** Same-unit N- and P-spikes unit occur in near synchrony. (**a**) The peak of the P-spike of a MM PYR (same as in **Aa**) occurs simultaneously with the N-spike trough. (**b**) The median time lag between N- and P-spikes in neocortical MM units is not consistently different from zero. Here and in **c**, ns, p>0.05, Wilcoxon’s test comparing to a zero null. Grey patch, 95% confidence limits. (**c**) The time lag between N- and P-spike in CA1 MM units is not consistently different from zero.

**Figure 2.**
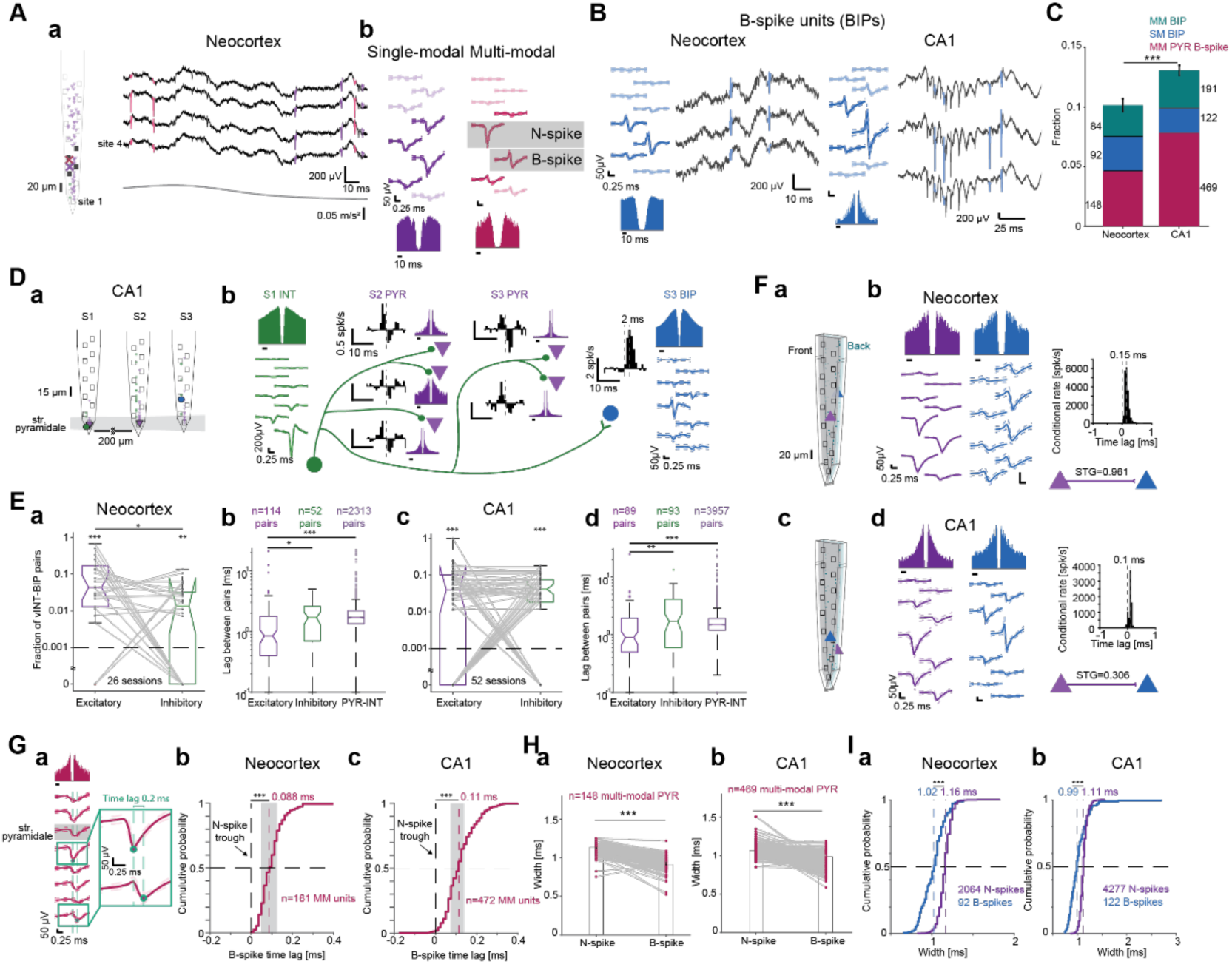
Biphasic extracellular spikes correspond to axonal potentials. **(A)** SM PYR and a MM unit with biphasic spikes (B-spikes) recorded from the neocortex. (**a**) **Left**, Schematic shank with 59 simultaneously-recorded units. (**b**) Wideband waveforms and ACHs. All conventions are the same as in **Fig. 1A**. **(B)** B-spikes appear without N-spikes. A biphasic unit (BIP) is defined as a unit with a dominant positive peak preceding a dominant negative peak in the main channel waveform. **(C)** The fraction of units with B-spikes is higher in CA1 than in neocortex. ***: p<0.001, G-test. **(D)** A BIP exhibiting “monosynaptic excitation” by a verified inhibitory interneuron (vINT). (**a**) Probe schematic in CA1. (**b**) Derived network of a vINT, BIP, five PYRs, and cross-correlation histograms (CCHs) of the vINT with the other units. The vINT is inhibitory for the five PYRs. However, the vINT-BIP CCH exhibits an excitatory monosynaptic peak. Note similar vINT and BIP ACHs. **(E)** (**a**) Fraction of vINT-BIP excitatory and vINT-BIP inhibitory connections in every session. Lined *: p<0.05, Wilcoxon’s test. Here and in **c**, dashed line, chance level; **/***: p<0.01/p<0.001, Binomial test comparing to chance level. (**b**) Time lags for vINT-BIP excitatory connections, vINT-BIP inhibitory connections, and PYR-INT excitatory connections. Here and in **d**, lined */**/***: p<0.05/p<0.01/p<0.001, U-test. (**c-d**) Same as **a-b**, for CA1. **(F)** (**a**) Neocortical PYR-BIP pair recorded from opposite sides of a 30 μm dual-sided probe. (**b**) Spike transmission gain (STG) is close to unity. Transmission peaks at a sub-millisecond, one-sided time lag. (**c-d**) PYR-BIP pair recorded using a dual-sided probe in CA1. **(G)** (**a**) Wideband waveforms and ACH of a MM PYR with a B-spike recorded from CA1. (**b-c**) Time lag of B-spike trough relative to same-unit N-spike trough. ***: p<0.001, Wilcoxon’s test compared to zero. All other conventions are the same as in **Fig. 1H**. **(H)** B-spikes are narrower than same-unit PYR N-spikes. ***: p<0.001, Wilcoxon’s test. **(I)** B-spikes of SM BIPs are narrower than N-spikes of SM PYRs or MM PYRs with P-spikes. ***: p<0.001, U-test.

Thus, non-negative spikes, which are not widely reported in the intact brain, may correspond to three possible compartmental sources of non-negative extracellular potentials during actional potentials. The axonal forward propagation, dendritic backpropagation, and dendritic return currents origins may be contrasted in freely-moving animals using high-density recordings by analyzing network activity. Sodium spikes generated at the axon initial segment (AIS) actively propagate forward along the axon and backwards into the soma and dendrites^37, 38^. Therefore, in extracellular recordings near the soma, potentials generated by axons and backpropagation into the dendrites are expected to exhibit sub-millisecond time lags with respect to somatic spikes due to conductance time. Axonal potentials are expected to be narrower than the somatic spikes due to higher density of voltage-gated potassium (Kv1) channels in the AIS^39^, whereas back-propagating spikes are expected to be wider due to lower dendritic density of K_A_ channels^40^. On the other hand, return currents in the proximal dendrites are expected to balance AIS spikes, and therefore be narrow and occur in synchrony with somatic spikes.

## Results

### Positive extracellular spikes correspond to return currents

To determine whether spikes of multiple polarities are observable in the intact brain, we recorded 9160 units from neocortex and hippocampal region CA1 of freely-moving mice (n=17; **Table S1**). Recordings were conducted using multi-shank arrays with vertical inter-electrode spacing of 15-20 μm (**Fig. 1Aa**).

Units with negative spike polarity were recorded by one or more electrodes on the same shank, denoted as “single-modal” (SM) units (**Fig. 1Ab**, **left**). However, we also recorded units with spikes of distinct polarities on different electrodes, denoted as “multi-modal” (MM) units (**Fig. 1Ab**, **right**). MM units were recorded in both neocortex (**Fig. 1A; Table S2**) and CA1 (**Fig. 1B; Table S3**). Overall, 966/9160 (10.5%) of the units were MM, with higher prevalence in CA1 (689/5971, 11.5%) compared with the neocortex (277/3189, 8.7%; p<0.001, G-test; **Fig. 1C**). Thus, in freely-moving mice, extracellular potentials of multiple polarities corresponding to a single spike are readily observable.

To determine whether extracellular P-spikes can be observed without N-spikes, we categorized the mean waveform on every electrode as an N-spike, a P-spike, a B-spike, or an undefined waveform according to the signal to noise ratio of the local extrema. Units for which the positive peak was larger than the trough on the electrode with the largest trough-to-peak magnitude were denoted P-spike units, “Punits”. We found SM Punits without N-spikes in both neocortex and in CA1 (**Fig. 1D**). The fraction of units with P-spikes was higher in neocortex (205/3189, 6.4%) compared with CA1 (207/5971, 3.4%; p<0.001, G-test; **Fig. 1E**). Thus, P-spikes may be observed not only together with N-spikes, but also in isolation.

The spatial dispersion of neocortical MM units with P-spikes was 85 [60 140] μm (median [interquartile interval, IQR]; n=15), compared to a dispersion of 75 [60 100] μm of SM pyramidal cells (PYRs; n=2049; p=0.027, U-test; **Fig. S1Aa**). In contrast, the dispersion of SM Punits was 40 [20 60] μm, smaller than of SM PYRs (p<0.001) or interneurons (INTs; 60 [40 100] μm; p<0.001, Kruskal-Wallis test corrected for multiple comparisons; **Fig. S1Ba**). Punit waveform magnitude was 81 [56 126] μV, smaller than PYR (116 [76 187] μV) or INT (100 [66 154] μV; p<0.001; **Fig. S1Ca**). Punits were not recorded closer to the edge of the probes, compared with PYRs or INTs (**Fig. S1Da**). The distance between same-unit N-and P-spikes was 20 [-30 30] μm (n=28 MM units; **Fig. S1Ea**). Similar results were observed in CA1 (**Fig. S1A-E**). Thus, MM units appear to capture extracellular potentials corresponding to multiple neuronal compartments during a spike.

Backpropagation into dendrites generates wider intracellular waveforms compared with somatic spikes^41^. To directly assess whether extracellular P-spikes correspond to backpropagation, we measured N- and P-spike widths using the inverse of the dominant frequency of the waveform spectrum. P-spike widths were 0.91 [0.8 1.04] ms, whereas same-unit N-spikes were wider at 1.14 [1.09 1.16] ms (n=25 neocortical MM units; p=0.0011, Wilcoxon’s test; **Fig. 1Fa**). Similar results were observed in CA1 (n=20 MM units; P-spikes: 1.11 [0.85 1.22] ms; N-spikes: 1.16 [1.07 1.28] ms; p=0.023; **Fig. 1Fb**). Comparing the widths of N- and P-spike waveforms regardless of the source unit, neocortical P-spikes widths were 1.02 [0.9 1.11] ms (n=179 SM Punits), whereas N-spikes were wider (1.16 [1.11 1.19] ms; n=2197 units; p<0.001, U-test; **Fig. 1Ga**). Similar results were observed in CA1 (P-spikes: 0.99 [0.88 1.11] ms; n=186 SM Punits; N-spikes: 1.11 [1.07 1.16] ms; n=4727 units; p<0.001, U-test; **Fig. 1Gb**). Thus, P-spikes recorded in near proximity of N-spikes are unlikely to be produced by backpropagation of somatic spikes into the dendrites.

To contrast the possibility that P-spikes recorded in close proximity to the N-spike correspond to axonal potentials with the possibility of dendritic return currents, we measured time lags between the trough of N-spikes and the peak of same-unit P-spikes (**Fig. 1Ha**). The median [95% confidence limits] time lag between N- and P-spike in neocortical MM units was -13 [-19 0] μs, not consistently different from zero (n=28 MM units; p=0.91, Wilcoxon’s test), and with an absolute value smaller than 19 μs (p=0.042, Wilcoxon’s test; **Fig. 1Hb**). Time lags in control pairs, consisting of two same-unit N-spikes with matched inter-electrode distances, were not consistently different (median [range]: 0 [-100,175] μs; n=7; p=1, Wilcoxon’s test). Similar results were observed in CA1, with the median [95%] time lags being 38 [25 38] μs (n=25; p=0.08) and absolute time lag smaller than 50 μs (p=0.041; **Fig. 1Hc**). Together, the waveform width and time lag analyses suggest that extracellular P-spikes that occur in close proximity to the soma correspond to dendritic return currents.

### Biphasic extracellular spikes correspond to axonal potentials

Action potential propagation in axonal and dendritic compartments was reported to generate triphasic spikes^18, 28, 32, 36^, but filtering with a high-pass frequency of 300 Hz transforms B-spikes into triphasic spikes (**Fig. S2A**). Because we used a very low cutoff frequency (0.1 Hz) and detrended the individual spikes, previously reported triphasic spikes likely correspond to B-spikes reported here. We recorded MM units with B-spikes in both neocortex (**Fig. 2A**) and CA1 (**Fig. S2B**).

Every spike was quantified using a biphasic index (BPI) which is -1 for a pure N-spike, 1 for a pure P-spike, and 0 for a symmetric B-spike (**Fig. S2C-E**). Units for which the electrode with the largest trough-to-peak magnitude contained a B-spike are denoted as B-spike units, “BIP” (**Fig. 2B**). The fraction of B-spike units was higher in CA1 (782/5971, 13.1%) than in neocortex (324/3189, 10.2%; p<0.001, G-test; **Fig. 2C**). Compared with P-spikes, B-spikes were more likely to be recorded with N-spikes (892 vs. 47 units; p<0.001, G-test, **Fig. 1E** and **Fig. 2C**). Thus, B-spikes are observable in extracellular recordings of freely-moving mice.

Although some violations exist^42–45^, most neurons use the same neurotransmitter on all postsynaptic targets^46^. If distinct compartments of a purely-inhibitory neuron are recorded as two distinct units, the unit representing somatic potentials will appear to excite the unit representing non-somatic potentials. Cross-correlation histogram (CCH) analysis identified a CA1 network with seven units that included an INT that made inhibitory monosynaptic connections with five PYRs (**Fig. 2Da**). The verified inhibitory INT (vINT) made an apparently excitatory monosynaptic connection with a BIP (**Fig. 2Db**). The auto-correlation histograms (ACHs) of the vINT and BIP had a rank correlation coefficient (cc) of 0.9 (p<0.001, permutation test), whereas the median [range] cc’s between the ACHs of the vINT and the PYRs were 0.2 [-0.15,0.8]. The apparent contradiction of Dale’s law is resolved if the BIP corresponds to non-somatic spikes of the vINT.

vINTs and BIPs were recorded simultaneously during 26/80 (32%) of the neocortical recording sessions, yielding a total of 2495 vINT-BIP pairs of which 166 were connected (6.7%). The number of excitatory connections (114/2495 pairs, 4.6%) was above chance (0.1%; p<0.001, Binomial test) and higher than the number of inhibitory connections (52/2495, 2.1%; p<0.001, G-test). The fraction of excitatory vINT-BIP pairs per session was 4.3% [1.1% 17%], higher than inhibitory vINT-BIP pairs (1.4% [0% 3.2%]; n=26 sessions; p=0.016, Wilcoxon’s test; **Fig. 2Ea**). Moreover, time lags for “excitatory” vINT-BIP connections were 0.8 [0.4 1.8] ms (n=114 pairs), shorter than for inhibitory vINT-BIP connections (1.7 [0.6 2.6] ms; n=52; p=0.019) and for excitatory PYR-INT pairs (1.7 [1.3 2.2] ms; n=2313; p<0.001, U-test; **Fig. 2Eb**). Similar results were observed in CA1, where 52/126 (41%) of the recording sessions had vINT-BIP pairs, with 182/2208 (8.2%) connected and 89/2208 (4%) excitatory pairs (p<0.001, Binomial test; **Fig. 2Ecd**). Thus, the network connectivity analysis suggests that SM BIPs correspond to non-somatic potentials.

Independent support for non-somatic origin of B-spikes comes from rare occasions in which two units corresponding to the same neuron were recorded simultaneously. In recordings from opposite sides of the same dual-sided shank (thickness, 30 μm; **Fig. 2Fa**), we found a neocortical PYR-BIP pair with near-unity spike transmission gain (STG) and a CCH peak at a sub-millisecond, unidirectional time lag (**Fig. 2Fb**). We found a similar setting in CA1 (**Fig. 2Fcd**). The high gain and short time lag are inconsistent with synaptic transmission in freely-moving mice^47^. Thus, the network vINT-BIP and the dual-sided PYR-BIP analyses suggest that B-spikes correspond to recordings from a non-somatic compartment.

The dendritic backpropagation and axonal potentials hypotheses both predict that B-spikes will occur after initial spike generation. B-spike troughs followed the N-spike troughs (**Fig. 2Ga**) in 161/161 (100%) of the neocortical MM units, with median [95% confidence limits] lags of 88 [75 100] μs (p<0.001, Wilcoxon’s test; **Fig. 2Gb**). Similar results were observed in CA1, where B-spikes occurred after the N-spikes in 466/472 (99%) of the cases (110 [100 110] μs; p<0.001; **Fig. 2Gc**). Thus, the trough time lag analysis is inconsistent with dendritic return currents, and is in agreement with both backpropagation and axonal potentials.

In brain slices, dendritic backpropagating spikes are wider than somatic spikes^41^ whereas axonal spikes are narrower^39^. Neocortical B-spikes had a width of 0.91 [0.87 0.98] ms, narrower than same-unit N-spikes (1.14 [1.09 1.16] ms; n=149 MM units; p<0.001, Wilcoxon’s test; **Fig. 2Ha**). Similar results were observed in CA1 (B-spikes: 0.98 ms [0.91 1.02] ms; N-spikes: 1.07 ms [1.02 1.11] ms; n=469; p<0.001; **Fig. 2Hb**). Comparing the widths of B- and N-spikes regardless of the source unit, we found that neocortical B-spikes widths were 0.88 [0.78 0.98] ms (n=92 BIPs), whereas N-spikes were wider (1.16 [1.11 1.19] ms; n=2064 units; p<0.001, U-test; **Fig. 2Ia**). Similar results were observed in CA1 (B-spikes: 0.97 [0.9 1.02] ms; n=122 SM BIPs; N-spikes: 1.11 [1.07 1.16] ms; n=4277 units; p<0.001; **Fig. 2Ib**). Thus, B-spikes recorded in near proximity to N-spikes are unlikely to represent backpropagation into the dendrites. Since both dendritic return currents and backpropagation are unlikely sources of B-spikes, axonal potentials are most likely.

### Positive phases of biphasic extracellular spikes correspond to return currents

The analysis of same-unit B- to N-spike time lag depended on the B-spike trough (**Fig. 2G**). Because by definition the peak of a B-spike occurs before the trough, we also measured the time lag of the B-spike peak relative to the same-unit N-spike trough (**Fig. 3Aa**).

**Figure 3.**
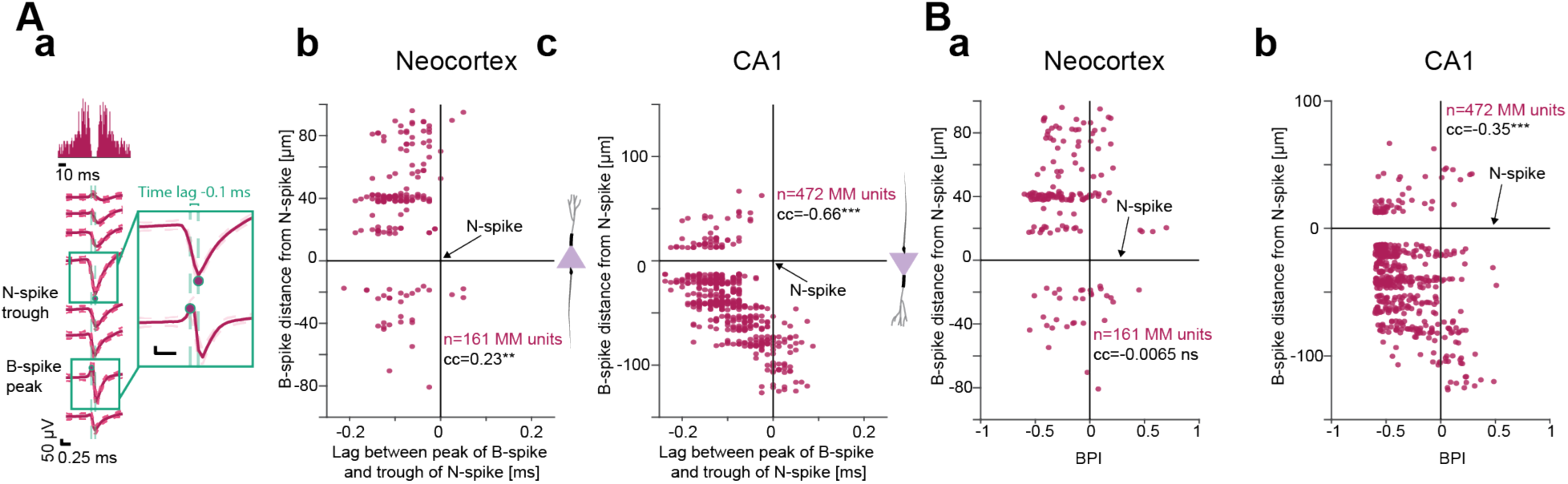
Positive phases of biphasic extracellular spikes correspond to return currents **(A)** (**a**) Computation of B-spike peak lag relative to the trough of the N-spike in the same MM unit. (**b-c**) Distance of B-spike from N-spike vs. lag of B-spike peak from N-spike trough. Distances are positive when the B-spike is closer to the surface of the brain. Cartoons illustrate the orientation of PYR soma, dendritic tree, and axon. Here and in **B**, cc, rank correlation coefficient; **/***: p<0.01/p<0.001, permutation test. The negative correlation in the neocortex (and the positive correlation in CA1) are consistent with the B-spike peaks near the soma and most proximal dendrites representing return currents during AIS spikes, and B-spike peaks near more distal dendrites representing return currents during somatic spikes. **(B)** Biphasic index (BPI) vs. distance between B- and N-spikes. The increasingly-positive BPIs farther from the N-spike (significant in CA1) are consistent with return currents forming a wavefront that propagates in space before the N-spike.

The median time lag was -88 μs (95% confidence limits: [-100 -75] μs; n=161 neocortical MM units; p<0.001, Wilcoxon’s test; **Fig. S4Aa**). Similar results were observed in CA1 (−100 [-113 -88] μs; n=472; p<0.001; **Fig. S4Ab**). Because N-spikes, B-spikes, and P-spikes form a continuum (**Fig. S2C-E**), a dichotomy between near-somatic P-spikes representing dendritic return currents and near-somatic B-spikes representing axonal potentials may be an oversimplification.

To determine the compartmental origin of the positive phase of B-spikes using blind extracellular recordings, we took advantage of the opposite axo-dendritic orientation of neocortical and CA1 PYRs (**Fig. 3Abc**, **insets**). We hypothesized that if there is a range of compartmental origins, the time lag between B-spike peak and N-spike trough will depend on the distance from the N-spike, and the correlation will be opposite in neocortex and CA1. Consistent with the prediction, in neocortex the cc was 0.23 (n=161 MM units with B-spikes; p=0.005, permutation test; **Fig. 3Ab**), whereas in CA1 the cc was -0.66 (n=472; p<0.001; **Fig. 3Ac**). Negative correlation was also observed when the distance was measured from the B-spike to the center of CA1 str. pyramidale (−0.36; p<0.001; **Fig. S4B**). Thus, the time lag vs. distance analyses indicate that B-spike peaks may either precede or be synchronized with same-unit N-spike troughs, suggesting that the B-spike peaks represent return currents.

For a spike generate at the AIS, return currents are expected at the proximal dendrites and the axon^15, 21^. However, once the spike begins propagating along the axon, proximal return currents are expected to be diminished due to refractoriness of voltage dependent channels, and the axonal return currents will be increasingly asymmetrical in space as the distance. Measuring the BPI of B-spikes in MM units as a function of the distance from the N-spike, we found that in neocortex the cc was 0.048 (n=161 MM units; p=0.56, permutation test; **Fig. 3Ba**), and in CA1 the cc was 0.42 (p<0.001; **Fig. 3Bb**). The polarity vs. distance results in CA1 suggest that spikes propagating along the unmyelinated part of the axon in close proximity with the N-spike exhibit an increasingly larger positive phase.

### BIPs and Punits exhibit place fields and phase precession

To directly examine whether Punits and BIPs are part of the CA1 local circuit, we compared spatial coding properties of PYRs, INTs, Punits, and BIPs while mice ran back and forth along a 150 cm linear track. Units of all four types exhibited increased firing rates at specific parts of the track (**Fig. 4A**).

**Figure 4.**
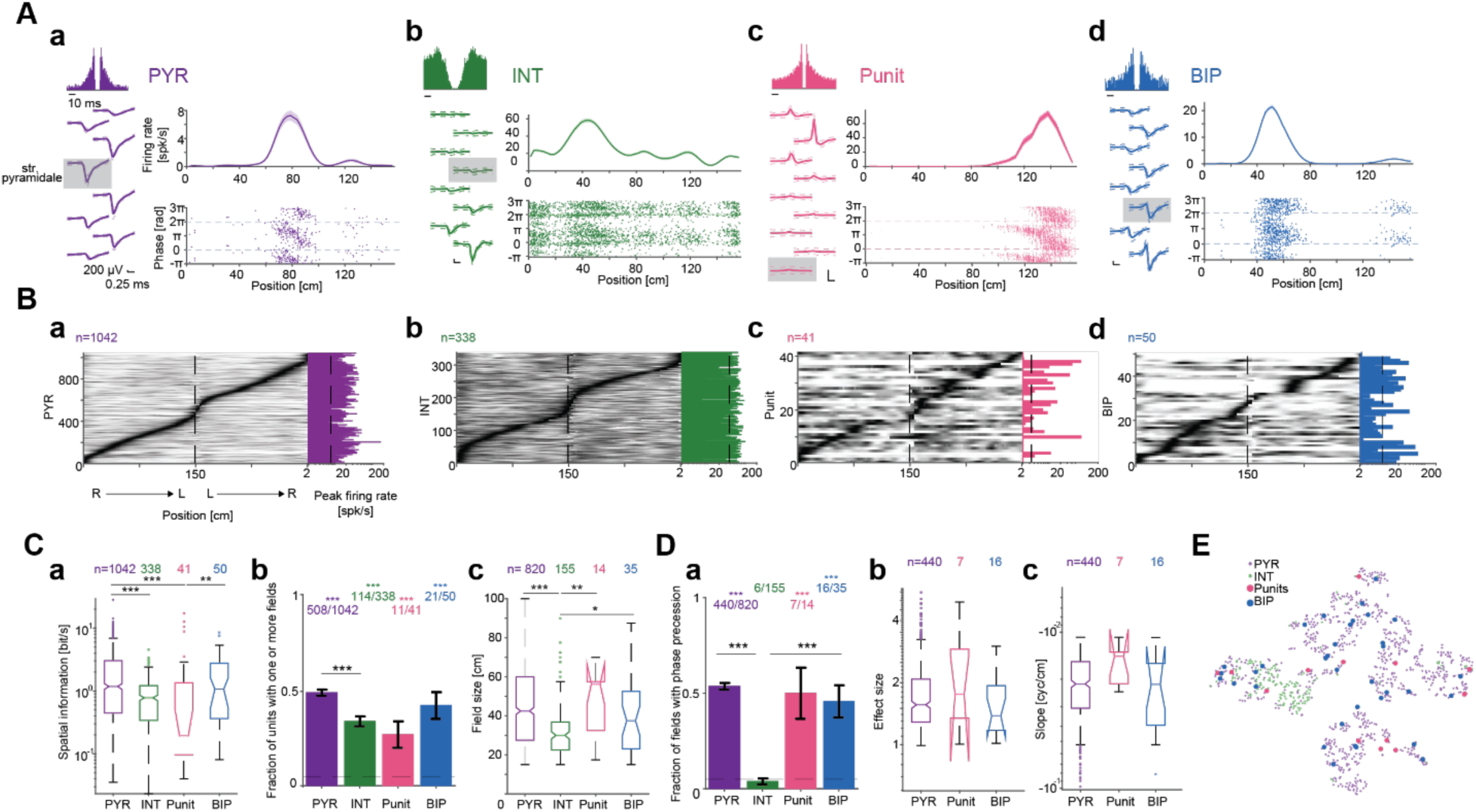
BIPs and Punits exhibit place fields and phase precession **(A)** Units recorded in CA1 as mice ran back and forth on a 150 cm long linear track. In every panel: **Left**, Wideband waveforms and ACH. **Top**, Firing rate as a function of position (mean±SEM over 82/96/70/41 same-direction trials of PYR/INT/Punit/BIP). **Bottom**, Instantaneous theta phase of every spike. Phase 0 corresponds to theta peak. **(B)** Units of all four types exhibit increased firing rates at specific regions of the linear track. (**a-d**) Every row represents a unit. Firing rates on right (R) to left (L) runs are concatenated with L to R runs and scaled to the 0 (white) to 1 (black) range. **(C)** Spatial rate coding of CA1 units. (**a**) Spatial information carried by the units. Top, number of units active and stable on the track. Here and in **c**, */**/***: p<0.05/p<0.01/p<0.001, Kruskal-Wallis test, corrected for multiple comparisons. (**b**) Fraction of units with one or more place fields. Here and in **Da**, ***: p<0.001, Binomial test, comparing to chance level of 0.05. Lined ***: p<0.001, G-test. Error bars, SEM. (**c**) Place field sizes. **(D)** Spatial phase coding of CA1 units. (**a**) Fraction of fields with theta phase precession. (**b**) Phase precession effect sizes. (**c**) Phase precession slopes. **(E)** t-SNE projection of all fields. Sample size is the same as in **Cc**. Features includes are spatial information, field size, in-field gain, TPP effect size, and TPP slope.

Focusing on active and stable units (**Fig. 4B**; **Table S4**), the spatial information carried by BIPs was 1.07 [0.35 2.76] bit/s (n=50), not consistently different from PYRs (1.18 [0.44 3.03] bit/s; n=1042; p=0.8; Kruskal-Wallis test; **Fig. 4Ca**). In contrast, Punits (0.19 [0.096 1.29] bit/s; n=41), and INTs (0.78 [0.34 1.22] bit/s; n=338) carried less spatial information than PYRs (p<0.001). 508/1042 (49%) of the PYRs, 114/338 (34%) of the INTs, 11/41 (27%) of the Punits, and 21/50 (42%) of the BIPs had one or more place fields (**Fig. 4Cb**). On average, BIPs had 1.67 fields/unit, not consistently different from PYRs (1.61; p=0.91, G-test; **Fig. S5A**). In-field gain was 9.5 [1.1 239.5] for BIPs, not consistently different from PYRs (5 [1.9 27.6]; p=0.92; Kruskal-Wallis test; **Fig. S5B**). Field sizes of Punits were 56.3 [30 57.5] cm (n=14 fields), not consistently different from PYR fields (42.5 [27.5 60] cm; n=820; p=0.78) or from BIP fields (37.5 [22.5 52.5] cm; n=35; p=0.56, Kruskal-Wallis test; **Fig. 4Cc**). Thus, in CA1, the spatial rate coding properties of BIPs are different from INTs but not distinct from PYRs.

As the animal advances through a place field, spikes of CA1 PYRs occur at progressively earlier theta phases, exhibiting theta phase precession^13, 48^. Precession was more prevalent among PYR fields (440/820, 53.7%) compared to INT fields (6/155, 3.9%; p<0.001), but not when compared with BIP (16/35, 45.7%; p=0.6) or Punit fields (7/14, 50%; p=0.88, G-test; **Fig. 4Da**). Among phase precessing PYR fields, precession effect size was 1.56 [1.29 2.07] (n=440), not consistently different from BIP (1.38 [1.17 1.8]; n=16; p=0.56) or Punit fields (1.76 [1 3.23]; n=7; p=0.94, Kruskal-Wallis test; **Fig. 4Db**). Finally, the slope of phase precessing PYR fields was -0.02 [-0.031 -0.016] cyc/cm, not consistently different from BIP (−0.02 [-0.042 - 0.019] cyc/cm; p=0.96) or Punit fields (−0.014 [-0.024 -0.013] cyc/cm; p=0.27, Kruskal-Wallis test; **Fig. 4Dc**). Thus, spatial phase coding of Punits and BIPs are not distinct from CA1 PYRs.

A t-SNE projection based on five spatial coding features (spatial information, field size, in-field gain, precession effect size, and precession slope) showed that BIP and Punit fields are interspersed within PYR fields (**Fig. 4E**). In summary, the similarity between the spatial coding properties of PYRs, Punits, and BIPs is consistent with non-negative potentials representing cellular compartments of local circuit CA1 PYRs.

### In CA1, Punits and BIPs occur outside str. pyramidale

In simultaneous recordings of neocortex and CA1, SM Punits were observed 200-300 μm above CA1 str. pyramidale, corresponding to the white matter in the corpus callosum (**Fig. 5A**).

**Figure 5.**
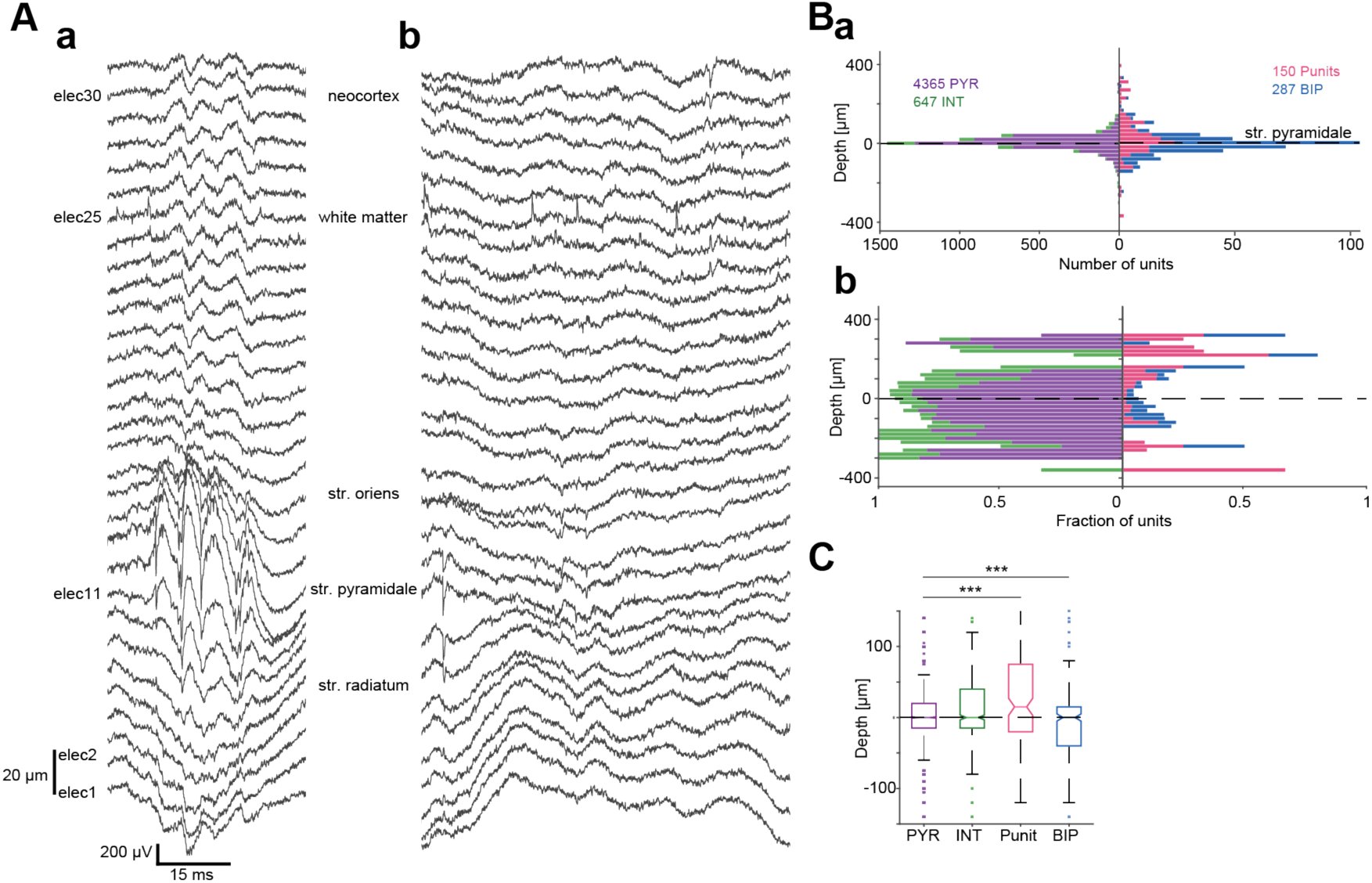
In CA1, Punits and BIPs occur outside str. Pyramidale. **(A)** (**a**) Wideband traces recorded by a 32-electrode array in a freely-moving mouse. Vertical inter-electrode spacing is 20 µm. A ripple event is evident, corresponding to str. pyramidale centered at elec11. (**b**) Traces from the same session showing multiple spike events, including a PYR in str. pyramidale (elec8-11), a SM Punit in the white matter (elec25), and a neocortical PYR (elec30-31). **(B)** (**a**) Depth distribution of PYRs, INTs, Punits and BIPs in CA1, relative to CA1 str. pyramidale. Bin size is 20 µm. Here and in **C**, horizontal dashed lines indicate the center of the layer. (**b**) Same as **a**, with absolute numbers scaled to fractions at every depth. **(C)** The depth distribution of non-negative units is distinct from the depth distribution of PYRs in CA1. ***: p<0.001, Kolmogorov-Smirnov test.

We determined which electrodes were located at the center of str. pyramidale based on spontaneous high frequency oscillation “ripple” events (**Fig. 5B**). In probes that span a vertical range of 300-620 μm, Punits appeared mainly in the white matter, 210 [-40 260] μm (n=30) above str. pyramidale. Most white matter Punits were SM units (29/30, 97%). Thus, consistent with reports from cat V1^23^, SM non-negative units correspond to axonal potentials in myelinated fibers.

Closer to str. pyramidale, the fraction of non-negative units was lowest at the center of the layer and higher in the flanking strata (**Fig. 5Bb**). The depth distribution of SM Punits (n=150) and SM BIP (n=287) differed from the depth distribution of SM PYRs (n=4365; p<0.001, Kolmogorov-Smirnov test; **Fig. 5C**). Whereas the median [IQR] depth of PYRs was 0 [-15 20] μm, Punits were closer to str. oriens (15 [-20 75] μm; p<0.001, U-test). Furthermore, the dispersion of Punit depths (SD, 115 μm) was higher than of PYRs (50 μm; p=0.004, permutation test). BIPs were recorded at depths of 0 [-40 15] μm, closer to str. radiatum compared to PYRs (p<0.001, U-test). Thus, the CA1 sub-layer organization of Punits and BIPs differs from PYRs, implying that the origin of Punits and BIPs is not somatic.

### Punits and BIPs precede PYRs and INTs in ripple lock in CA1

As an independent assessment of the possibility that Punits and BIPs represent non-somatic potentials of local circuit neurons in CA1, we analyzed spiking activity during ripple oscillations. Akin to PYR and INT, firing rates of non-negative units increased during ripples (**Fig. 6A**).

**Figure 6.**
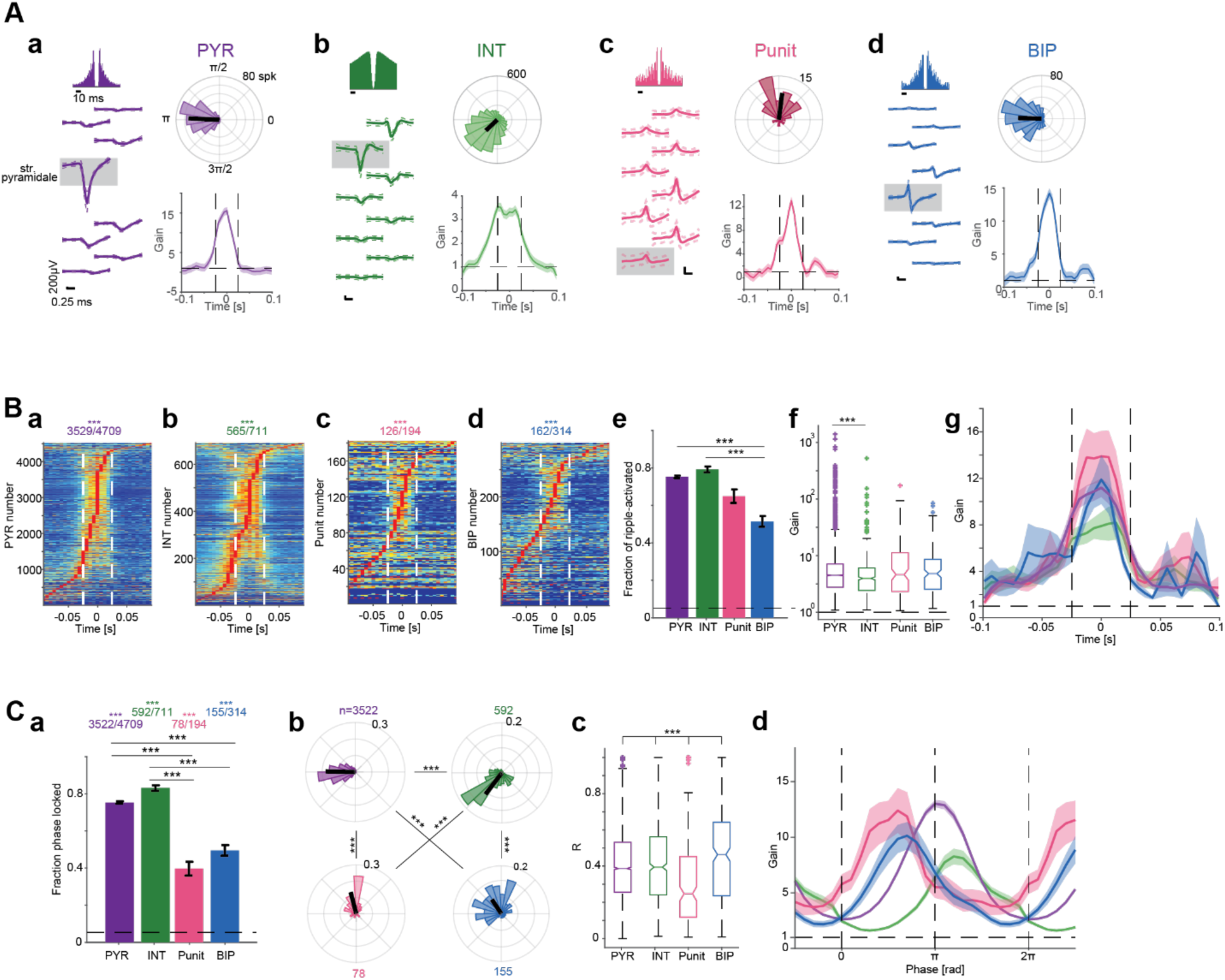
Punits and BIPs precede PYRs and INTs in ripple lock in CA1. **(A)** Units with non-negative waveforms exhibit time-locking to ripple events. (**a-d**) Example ripple-locked CA1 PYR, INT, Punit, and BIP. **Left**, Wideband spike waveforms and ACHs. **Top right**, Ripple phase during every spike, binned into 20 equal-sized bins. Black line represents the mean phase and the resultant length. **Bottom right**, Firing rate gain as a function of time in ripple. Here and in **B**, vertical dashed lines indicate mean ripple time range. **(B)** Punits and BIPs exhibit increased firing rates during ripple events. (**a-d**) Ripple-triggered firing rate histograms of all CA1 units. For presentation purposes, firing rates are scaled to the 0 (blue) to 1 (red) range. ***: p<0.001, Binomial test comparing to chance level, 0.05. (**e**) The fraction of BIPs with firing rate gain above 1 during ripple events. Here and in **Ca**, lined ***: p<0.001, G-test, corrected for multiple comparisons. Error bars, SEM. (**f**) Firing rate gain during ripples. Here and in **Cc**, ***: p<0.001, Kruskal-Wallis test. (**g**) Gain as a function of time in ripple. **(C)** Phase-locked Punits and BIPs precede PYRs and INTs in ripple events. (**a**) The fraction of phase-locked units. (**b**) Punits and BIPs spike earlier on the ripple cycle compared to PYRs and INTs. ***: p<0.001, Wheeler-Watson test. (**c**) Resultant lengths of ripple phase, indicating lock strength. (**d**) Gain as a function of phase in ripple.

Units of all four populations exhibited consistent (p<0.05, Poisson test) firing rate increases during ripple events (p<0.001, Binomial test; **Fig. 6Ba-d**). The fraction of BIP with increased ripple firing rates was 162/314 (51.6%), lower than PYRs (3529/4709, 75%; p<0.001) and INT (565/711, 79.5%; p<0.001), but not consistently different from Punits (126/194, 65%; p=0.75, G-test; **Fig. 6Be**). Of the units with increased ripple firing rates, the gain of Punits (4.8 [2.5 8.6], n=126) and BIP (4.62 [2.12 11.28], n=162) was not consistently different from PYRs (4.47 [2.77 7.13], n=3529; p=0.96 and p=0.89, respectively; Kruskal-Wallis test; **Fig. 6Bf**). Thus, the firing rates of Punits and BIPs are temporally modulated by ripples (**Fig. 6Bg**), consistent with spiking of CA1 local circuit neurons.

The fraction of phase-locked Punits (78/194, 40.2%) and BIPs (155/314, 49.4%) was above chance (p<0.001, Binomial test; **Fig. 6Ca**) but lower than phase-locked PYRs (3522/4709, 74.8%) and INTs (592/711, 83.3%; p<0.001, G-test; **Fig. 6Ca**). Due to the curvature of dorsal CA1, a parallel electrode array inserted perpendicularly to the point of penetration may record only non-somatic compartments of some neurons (**Fig. 6Acd**). Superficial units (closer to str. radiatum) spike earlier in the ripple cycle than deep units (closer to str. oriens^49^). Every population fired at a different phase: earliest were Punits (0.53π [0.46π 0.87π]; n=78), followed by BIPs (0.65π [0.46π 1.02π]; n=155), PYRs (1.01π [0.89π 1.11π]; n=3522) and finally INTs (1.25π [1.11π 1.48π]; n=592; p<0.001, Wheeler-Watson test; **Fig. 6Cb**). The resultant length R, indicating the strength of phase lock, of Punits (0.35 [0.14 0.53]) was lower than of PYRs (0.42 [0.3 0.54]) and INT (0.42 [0.26 0.57]), and highest for BIP (0.55 [0.39 0.67]; p<0.001, Kruskal-Wallis test; **Fig. 6Cc**). Thus, Punits and BIPs are phase-locked to high frequency ripples but spike earlier than PYRs in the ripple cycle, suggesting that Punits and BIPs correspond to PYR somata that reside in superficial CA1 layers.

## Discussion

Using high density extracellular recordings from freely-moving mice, we investigated the possible origin of non-negative extracellular waveforms. We found that P-spikes and positive phases of B-spikes are most consistent with return currents, and that isolated P- or B-spikes are most consistent with axonal potentials. In hippocampal region CA1, Punits and BIPs carry spatial information, exhibit phase precession, and are phase locked to ripples, akin to local circuit PYRs.

### Compartmental origin of non-negative extracellular spikes

It was suggested that P-spikes in vitro^17^ and B-spikes in vivo^34, 35^ correspond to active backpropagation of somatic spikes into dendrites^41^. Previously-reported non-negative spikes were seen far from the somatic potentials^36^, whereas our P- and B-spikes were in close proximity to the N-spikes. Backpropagating spikes are wider than somatic signals^40, 41^. Because the P- and B-spikes in the present dataset are narrower than N-spikes, the non-negative spikes are unlikely to be produced by backpropagation.

Return currents represent the instantaneous outward flow of positive ions during an action potential, balancing charges and maintaining a steady membrane potential. P-spikes in vitro^19, 20^ and in silico^15, 16^ were suggested to correspond to return currents. Up to charge movement times, return currents should occur at the same time as spike generation at the AIS^50^. Therefore, spikes representing return currents are expected to occur in-sync or slightly before the somatic spike. We found that P-spikes occur in synchrony with same-unit N-spikes, and that the positive phase of B-spikes occurs before the same-unit N-spikes, supporting the interpretation that positive spike phases represent return currents.

B-spikes in vitro^7, 24–27^, in silico^9, 15, 21^, and in vivo^22, 28, 29^, and P-spikes^22, 23^ were suggested to correspond to axonal potentials. The positive peak of a B-spike far from the N-spike can be viewed as a local return current. As an AIS-generated spike propagates along the axon, the return currents propagate as well. For an unmyelinated juxta-somatic axon, the return currents of the propagating spike are expected distally because the sodium channels proximal to the soma are refractory. When recorded at a single point in space near the axon over time, the wavefront of the positive peak occurs before the negative spike, yielding a peak-then-trough biphasic spike.

Sodium spikes can be generated not only in the soma/AIS but also in the dendrites^51–53^. In vitro^18, 32^ and in silico^21, 33^, B-spikes were suggested to be generated in the dendrites by forward propagation of dendritic sodium spikes. In MM units, P-spikes were in sync with N-spikes, and the trough of the B-spikes occurred after the N-spike trough. Therefore, it is unlikely that P- and B-spikes were created by forward propagation of dendritic sodium spikes. Dendrites can also generate calcium spikes^54, 55^. In a dendritic calcium spike, P- and B-spikes are expected to appear before the somatic spike, but calcium spikes are much wider. Thus, the most likely compartmental counterparts of MM P- and B-spikes are dendritic return currents and axonal potentials.

Isolated P- and B-spikes both corresponded to axonal spikes. Return currents in myelinated axons can flow via nodes of Ranvier, manifesting as isolated positive phase potentials that correspond to SM Punits. It is possible that the difference between the P- and B-spike waveforms originates from difference in the myelin sheath. In neocortical PYRs, the axon becomes myelinated 35-50 μm after the AIS^56^, 75-90 μm from the soma^57^. INT axons myelinate closer to the soma, 20-50 μm^58^. Axons of CA1 PYRs myelinate even closer, within 10-20 μm^59^. The early myelination for neocortical INTs and CA1 PYRs may explain the scarcity of MM INTs and the lower prevalence of MM units in CA1.

### Alternative, non-compartmental sources of non-negative spikes

The short time-lags of excitatory vINT-BIP pairs and the high-fidelity PYR-BIP transmission may suggest that B-spikes are generated by inter-neuronal gap junctions^60–62^. However, gap junction prevalence among neocortical PYRs is low^62^, inconsistent with the observation of neocortical MM PYRs with non-negative spikes. Moreover, neurons coupled by gap junctions typically yield CCHs with two-sided peaks^61, 63^, whereas the CCHs of the excitatory vINT-BIP and PYR-BIP pairs are unidirectional. Therefore, it is unlikely that “excitatory” connections to BIP were caused by gap junctions.

A possible source of SM Punits is sporadic juxtacellular recording, established by electrode adhesion of to the membrane^64^. However, the sizes of the electrodes employed (10 x 15 μm) and the diameter of a PYR soma (10-15 μm^65^) are similar. Furthermore, SM Punit waveforms were recorded together with waveforms of other units by the same electrode. Thus, non-negative extracellular waveforms are unlikely to be due to juxtacellular recordings.

In principle, P-spikes can be generated by somatic potassium action potentials. Potassium spikes were previously observed as slow “hyperpolarizing” responses in invertebrate muscles^66, 67^, and may also occur when extracellular potassium concentrations are very high or when potassium channels are inactivated^68^.

However, the presently-reported P-spikes occur together with N-spikes and are much sharper than previously-reported potassium spikes.

### Observability

Non-negative spikes comprise about 10% of the waveforms, but past studies rarely reported non-negative spikes. The discrepancy may be due to low-density recordings, limited detection parameters, and spike sorting bias. To minimize waveform distortion due to filtering^69, 70^, we employed a very low high-pass frequency (0.1 Hz). We expect the application of similar signal processing techniques to increase the prevalence of non-negative spikes reported in the future.

The fraction of P-spikes was higher in neocortex compared with CA1. The lower prevalence of P-spike units in CA1 may be due to a shadowing effect, caused by the high density of PYRs in str. pyramidale.

Due to the layered organization of hippocampal region CA1, the shanks of a probe implanted vertically in the dorsal hippocampus and centered on str. pyramidale are parallel to the axo-dendritic tree of the PYRs. Therefore, B- and N-spikes of CA1 PYRs may be recorded simultaneously, yielding MM PYRs, consistent with the higher probability of observing MM units with B-spikes in CA1 compared with the neocortex. In contrast, SM BIP and Punits are likely to represent compartments of units with unrecorded somata. Isolated compartmental recordings may represent somata outside str. pyramidale, or PYRs whose axo-dendritic axes are not parallel to the probe.

It is a-priori possible that some of the CA1 Punits and BIP represent non-somatic compartments of CA3 neurons projecting to CA1. Place coding in CA3 PYRs includes higher fraction of place fields, larger place fields, higher spatial information, and lower phase precession prevalence^71^. However, the prevalence and properties of place coding by SM CA1 BIPs and Punits are not consistently different from CA1 PYR place coding. Thus, the properties of Punits and BIPs recorded in CA1 are inconsistent with CA3 place coding. Because studies of hippocampal place coding typically analyze spike timing but spike waveforms are only rarely shown, it is unknown whether previous studies considered Punits and BIPs as place cells.

### Limitations

While the present dataset yielded many non-negative waveforms, the structure of the multi-shank probes (vertical span of 100-200 μm) and electrode size (10 x 15 μm) are optimized for high-density somatic recordings. The limited vertical span precludes monitoring distant potentials, and the prevalence of MM units may be even higher with higher-span probes^2^. A large electrode averages potentials from multiple small sources: the diameter of a neocortical PYR axon is 0.2-0.5 μm^72^, and the diameter of an apical proximal dendrite is 1-4 μm^73^. Smaller electrodes (e.g., 7 μm or smaller^74^) may capture more potentials generated by non-somatic compartments, with the possible caveat of higher impedance and thermal noise.

### Future directions

Punits and BIP were most consistent with return currents and axonal spikes, but we did not demonstrate causality. A causal link may be achieved in using compartmental blockade. Recording from the CA1 feedforward network, a specific blocker for axonal sodium channels (NaV1.2^75, 76^) may be applied locally. We predict that MM PYRs with B-spikes recorded before and during the blockade will turn to SM PYRs, and that BIPs will disappear. Alternatively, if sodium channels in proximal dendrites are blocked (NaV1.6^77^), the N-spike to P-spike distance is expected to increase in MM PYRs with P-spikes.

The analysis of P- and B-spikes may have direct implications for studying neural diseases. Presently, measurements of axonal propagation velocity are carried out ex vivo^58, 78–81^ or under anesthesia^82^. Future studies may measure axonal propagation velocity in the intact brain directly by computing time lags between N- and B-spikes in MM units with B-spikes. The approach may enable studying neurological disorders associated with axonal pathology, including Alzheimer’s disease^83^ and multiple sclerosis^84, 85^.

## Acknowledgements

We thank Jasmine Herszage and Lior Oppenheim for constructive comments. This work was supported by the United States-Israel Binational Science Foundation (BSF; 2015577); by the European Research Council (679253); by the Israel Science Foundation (638/16); by the Israel Science Foundation FIRST Program (1871/17); by the Rosetrees Trust (A1576); and by the Canadian Institutes of Health Research (CIHR), the International Development Research Centre (IDRC), the Israel Science Foundation (ISF), and the Azrieli Foundation (2558/18).

## Author contributions

S.S. and E.S. conceived and designed the study. S.S., A.L., H.E.S., L.S., R.G., and E.S. constructed optoelectronic probes and implanted animals. S.S., A.L., H.E.S., L.S., and R.G. carried out experiments. S.S., A.L., H.E.S., and E.S. analyzed data. S.S. and E.S. wrote the manuscript, with input from all authors.

## Materials and Methods

### Experimental animals

A total of 17 freely moving mice were used in this study (**Table S1**). 16 of the mice were males and one was female. Four of the mice were hybrid, the first-generation hybrid offspring (FVB/NJ x C57BL/6J)F1, and 13 were on a C57BL background^86^. The mice aged 8-32 weeks (median, 16.5 weeks) at the time of implantation. Animals were healthy, were not involved in previous procedures, and weighed 24.2-35.9 g (median, 30.5 g) at the time of implantation. Mice were single housed to prevent damage to the implanted apparatus. All animal handling procedures were in accordance with Directive 2010/63/EU of the European Parliament, complied with Israeli Animal Welfare Law (1994), and approved by Tel Aviv University Institutional Animal Care and Use Committee (IACUC #01-16-051).

### Probes and surgery

Every animal was implanted with a multi-shank silicon probe attached to a movable microdrive^87^. The probes used were Stark64 (Diagnostic Biochips; six mice), Buzaski32 (NeuroNexus; five mice), Linear32 (A1×32-Edge-10mm-20-177, NeuroNexus; four mice) and Dual-sided64 (DS64, Diagnostic Biochips; two mice). The Stark64 probe consists of six shanks, spaced horizontally 200 μm apart, with each shank consisting of 10-11 recording sites, spaced vertically 15 μm apart. The Buzaski32 probe consists of four shanks, spaced horizontally 200 μm apart, with each shank consisting of eight recording sites, spaced vertically 20 μm apart. The Linear32 probe consists of one shank, with 32 recording sites, spaced vertically 20 μm apart. The DS64 probe consists of two 30 μm thick dual-sided shanks, spaced horizontally 250 μm apart, with each shank consisting of 16 channels on each side (front and back), spaced vertically 20 μm apart. Probes were implanted in the neocortex above the right hippocampus (PA/LM, 1.6/1.1 mm; 45° angle to the midline) under isoflurane (1%) anesthesia as previously described^88^.

### Recording sessions

Recordings sessions lasted 4.3 [3.4 5.7] h (median [IQR] of 198 sessions). After every session, the probe was translated vertically downwards by no more than 70 μm. All hippocampal recordings were from the CA1 layer, recognized by the appearance of multiple high-amplitude units and iso-potential spontaneous ripple events. In every session, neural activity was recorded while the animal was in the home cage. Animals were equipped with a 3-axis accelerometer (ADXL-335, Analog Devices) for monitoring head movements.

### Linear track sessions

During every session that involved running on the linear track (**Table S4**), neural activity was first recorded while the animal was in the home cage for at least 40 min. The animal was then placed on a 150 cm linear track that extended between two 10 x 10 cm square platforms. Each platform included a water delivery port. Mice were under water restriction and were trained to repeatedly traverse the track for a water reward of 3-10 μl. Head position and orientation were tracked in real time using two head-mounted LEDs, a machine vision camera (ace 1300-1200uc, Basler), and a dedicated system (Spotter^89^). Four mice ran a total of 75 sessions that included 168 [130 212] one-direction trials over 50-90 min. Trials with a mean running speed below 10 cm/s were excluded from analyses.

### Spike detection and sorting

Neural activity was filtered, amplified, multiplexed, and digitized on the headstage (0.1–7500 Hz, x192; 16 bits, 20 kHz; RHD2132 or RHD2164, Intan Technologies), and then recorded by an RHD2000 evaluation board (Intan Technologies). Offline, spikes were detected, detrended, and sorted into single units automatically using KlustaKwik3^90^ for shanks with up to 11 sites/shank, or KiloSort2^91^ for shanks with 16-32 channels. Automatic spike sorting was followed by manual adjustment of the clusters. Only well-isolated units were used for further analyses (amplitude >40 μV; L-ratio <0.05; inter-spike interval index <0.2; ^47^).

### Categorization of spike waveforms

For every well-isolated unit, the mean waveform recorded on every electrode was categorized as an N-spike, P-spike, B-spike, or left uncategorized. Waveform categorization proceeded as follows. First, the waveforms of all spikes were averaged over all spikes, and the mean and SD were denoted. The mean of first 0.15 ms (3 samples) was subtracted from the mean waveform, and then the mean and the SD waveform were upsampled four-fold in time using cubic spline interpolation, yielding two 128-sample vectors. Second, all local extrema of the mean upsampled vector were detected. The local extremum with maximal value *p* was denoted as the “peak”, and the local extremum with the minimal value *n* was denoted as the “trough”. For every waveform, a bipolar index was computed, defined as BPI=(p-|n|)/(p+|n|). The BPI ranges -1 to 1, taking a value of zero when the peak and trough have the same absolute value (**Fig. S2C-E**).

B-spikes: If the peak preceded the trough, the peak was larger than 1.25 SDs at the peak and the trough was larger than one SD at the trough, and -0.6<BPI<0.8, the waveform was categorized as a B-spike.

P-spikes: A waveform not categorized as a B-spike that had a peak larger than the trough p>|n|, and the peak was larger than 1.75 SDs at the peak, was categorized as a P-spike.

N-spikes: A waveform not categorized as a B-spike that had a trough larger than the peak |n|>p, and an absolute value of the trough larger than 1.75 SDs at the trough, was categorized as an N-spike.

All other waveforms were uncategorized. Overall, we categorized 2112 spikes as B-spikes, 1434 as P-spikes, and 37255 as N-spikes. 60383 spikes were not categorized. A median [IQR] of 4 [3 6] spikes were categorized per unit. We verified that the categorization of waveforms was not sensitive to the specific parameter values noted above. In particular, when using a symmetric definition for B-spikes (one SD at the peak and one SD at the trough), and equivalent definitions for P-spikes and N-spikes (two SDs at the peak and trough, respectively), the number of B-/P-/N-spikes were 3123, 1097, and 32652.

### Classification of units

For every unit, the waveform with the maximal magnitude, defined as the difference between the maximal value and the minimal value, was denoted the “main channel”. Every unit was then classified according to the spike category of the waveform in the main channel. Thus, units with a B-spike in the main channel were classified as “B-spike units” (BIPs); units with a P-spike in the main channel were classified as “P-spike units” (Punits); and units with an N-spike in the main channel were classified as “N-spike units”.

In addition, every unit was classified as a single-modal (SM) or a multi-modal (MM) unit. If all categorized constituent spikes had the same categorization (e.g., all were N-spikes), the unit was classified as SM. All other units were classified as MM.

Finally, negative-spike units were classified into putative pyramidal cells (PYRs) or PV-like interneurons (INTs) using a Gaussian mixture model^92^.

A total of 9160 units were recorded from the 17 mice during 198 sessions (**Table S1**). Of these, 6959 (76%) were PYRs, 1334 (14.6%) were INTs, 378 (4.1%) were Punits, and 489 (5.3%) were BIPs. As for the categorization of spikes, we verified that the classification of units was not sensitive to the specific parameter values. In particular, when using the alternate set of parameter values, the number of PYR/INT/Punits/BIP were 6879/1323/375/571. A total of 3189 units were recorded from the neocortex during 78 sessions (**Table S2**), and 5971 units were recorded from CA1 during 126 sessions (**Table S3**).

### Determining monosynaptic connectivity

To determine whether a monosynaptic connection may exist between units, count cross-correlation histograms (CCHs; 0.1 ms bins) were constructed for putative pre- and postsynaptic spike train pairs as previously described^93^. Briefly, the spike transmission curve was estimated by the difference between the deconvolved CCH and the baseline, determined by hollowed median filtering of the count CCH, scaled to spikes/s. The spike transmission gain (STG) was defined as the area under the peak in the monosynaptic temporal region of interest (ROI; 0<t≤5 ms), extended until the causal zero-crossing points. Units that participated as a reference in a CCH that exhibited a consistent peak (p<0.001, Bonferroni-corrected Poisson test on the deconvolved count CCH compared to baseline) in the monosynaptic ROI were defined as presynaptic excitatory cells. Units that participated as a reference in a CCH that exhibited a consistent trough in the monosynaptic ROI were defined as presynaptic inhibitory cells.

### Identification of place fields

All analyses were performed only on “active and stable” units^94^. Briefly, “active” units fired a minimum of 5 spikes in at least one 2.5 cm spatial bin on the linear track, pooled over all trials. To determine whether a unit was “stable”, all rank correlation coefficients between the firing rate maps of same-direction trial pairs were computed. Statistical significance was determined by the geometric mean p-values derived from a permutation test over all pairs in both running directions. Units with a p-value below chance were considered stable. Overall, 1471/4515 (32.6%) of the recorded units were active (**Table S4**).

For every active and stable unit, place fields were defined in regions of space in which the on-track firing rate deviated from chance level, determined recursively based on a Poisson distribution^94^. Place fields that contained fewer than 30 spikes were omitted from further analyses. Precession effect size was quantified as the ratio between the fit of spikes to a circular-linear model divided by the median of 300 model fits to randomly permuted phase/position pairs^86^.

### Analysis of ripples

High frequency ripple oscillations in CA1 were detected as previously described^49^. Briefly, ripples were detected independently at each electrode. The wide-band signal was bandpass-filtered (80-250 Hz; difference-of-Gaussians, DOG; zero-lag, linear phase FIR), and instantaneous power was computed by clipping values above five SDs, rectifying, and low-pass filtering. The mean and SD were computed from the power of the clipped signal during non-theta immobility. Events for which the power of the unclipped signal exceeded five SDs from the mean were detected, expanded until the power fell below two SDs to define event edges, and aligned by the trough closest to the peak power. The center of the CA1 pyramidal cell layer was determined for each shank by the maximal ripple amplitude, and region assigning was done per unit. Subsequent analyses were based on ripples detected at the center of the layer. For every unit, ripple gain was defined by the ratio of the firing rate during spontaneous ripple events and the baseline firing rate of that unit, as measured during non-theta immobility. The instantaneous phase of the DOG-filtered signal was derived from its Hilbert transform, and spikes that occurred during a ripple event were assigned the phase (0-2π) at the time of firing.

### Statistical analyses

In all statistical tests a significance threshold of α=0.05 was used. An exception was the threshold used for determining whether two units exhibit monosynaptic connectivity (α=0.001). In all cases, non-parametric testing was used. An exception was to determine ripple phase locking (Rayleigh test). All statistical details (n, median, IQR, range, confidence limits, mean, SEM) can be found in the main text, figures, figure legends, and tables. To estimate whether fractions were larger or smaller than expected by chance, an exact one-tailed Binomial test was used. Differences in the proportions of two categorical variables were tested with a likelihood ratio (G-) test (two-tailed). Bonferroni’s correction was employed in cases of G-test multiple comparisons. Differences between two group medians were tested with either Mann-Whitney’s U-test (unpaired samples) or Wilcoxon’s paired signed-rank test (two-tailed). Differences between medians of four groups were tested with Kruskal-Wallis test, and corrected for multiple comparisons using Tukey’s procedure. Wilcoxon’s signed-rank test was employed to determine whether a group median is distinct from a predetermined value (two tailed). Association between parameters was quantified using Spearman’s rank correlation and tested with a permutation test. The statistical significance of unimodal phase locking was tested using Rayleigh’s likelihood-ratio test of uniformity. Differences between the circular medians of two groups were tested with Wheeler-Watson’s nonparametric two-sample test (two-tailed). For all figures, *: p<0.05; **: p<0.01; ***: p<0.001.

## Tables

**Table S1.** Mice used for electrophysiological recordings

| Animal ID | Sex | Age<sup>1</sup><br>[week] | Weight<sup>1</sup><br>[g] | Strain<sup>2</sup> | Probe | Sessions | Units<sup>3</sup> | Neocortex<br>sessions | CA1<br>sessions | CA1 linear<br>track<br>sessions |
| --- | --- | --- | --- | --- | --- | --- | --- | --- | --- | --- |
| mA234 | M | 16 | 30 | HYB | Buzsaki32 | 40 | 2285 | 14 | 26 | 24 |
| mB142 | M | 20 | 35.9 | HYB | Buzsaki32 | 1 | 10 | 1 | 0 | 0 |
| mC41 | M | 10 | 33.7 | HYB | Stark64 | 37 | 2473 | 5 | 32 | 32 |
| mDL5 | M | 32 | 30.9 | C57 | Buzsaki32 | 14 | 220 | 14 | 0 | 0 |
| mDS1 | M | 14 | 25.7 | C57 | DS64<sup>4</sup> | 21 | 861 | 4 | 17 | 5 |
| mDS2 | F | 30 | 24.2 | C57 | DS64<sup>4</sup> | 12 | 1546 | 9 | 3 | 0 |
| mF105 | M | 26 | 32.5 | HYB | Stark64 | 1 | 57 | 1 | 0 | 0 |
| mF108 | M | 12 | 31.4 | C57 | Stark64 | 2 | 138 | 2 | 0 | 0 |
| mF79 | M | 17 | 30.1 | C57 | Stark64 | 1 | 41 | 1 | 0 | 0 |
| mF84 | M | 8 | 24.4 | C57 | Linear32 | 8 | 126 | 8 | 0 | 0 |
| mF93 | M | 18 | 33.5 | C57 | Stark64 | 2 | 170 | 2 | 0 | 0 |
| mK01 | M | 24 | 29.5 | C57 | Linear32 | 5 | 67 | 3 | 5 | 0 |
| mO251 | M | 13 | 31.5 | C57 | Linear32 | 5 | 52 | 2 | 4 | 0 |
| mP101 | M | 16 | 29.7 | C57 | Buzsaki32 | 6 | 231 | 1 | 5 | 0 |
| mP20 | M | 20 | 31 | C57 | Stark64 | 4 | 165 | 4 | 0 | 0 |
| mP23 | M | 14 | 31.2 | C57 | Buzsaki32 | 26 | 666 | 3 | 23 | 14 |
| mV99 | M | N/A | 25.8 | C57 | Linear32 | 13 | 52 | 4 | 11 | 0 |
| **Total** |  |  |  |  |  | **198** | **9160** | **78** | **126** | **75** |

<sup>1</sup> At the time of implantation.
<sup>2</sup> HYB, hybrid mice. C57, mice on a C57BL background.
<sup>3</sup> Total number of well-isolated units recorded during every session.
<sup>4</sup> Dual-sided64 probe.

**Table S2.** Units recorded from neocortex

|  |  |  | MM |  |  |  |  | SM |  |  |  |  |
| --- | --- | --- | --- | --- | --- | --- | --- | --- | --- | --- | --- | --- |
| Animal ID | Sessions | Units | Total | PYR | INT | Punit | BIP | Total | PYR | INT | Punit | BIP |
| **mA234** | 14 | 586 | 37 | 15 | 4 | 2 | 16 | 549 | 360 | 137 | 35 | 17 |
| **mB142** | 1 | 10 | 2 | 1 | 0 | 0 | 1 | 8 | 5 | 1 | 1 | 1 |
| **mC41** | 5 | 147 | 3 | 0 | 0 | 0 | 3 | 144 | 101 | 12 | 22 | 9 |
| **mDL5** | 14 | 220 | 3 | 2 | 0 | 0 | 1 | 217 | 158 | 47 | 9 | 3 |
| **mDS1** | 4 | 58 | 1 | 1 | 0 | 0 | 0 | 57 | 27 | 21 | 5 | 4 |
| **mDS2** | 9 | 1351 | 179 | 126 | 11 | 5 | 37 | 1172 | 942 | 187 | 29 | 14 |
| **mF105** | 1 | 57 | 0 | 0 | 0 | 0 | 0 | 57 | 43 | 11 | 2 | 1 |
| **mF108** | 2 | 138 | 12 | 3 | 1 | 0 | 8 | 126 | 56 | 53 | 7 | 10 |
| **mF79** | 1 | 41 | 0 | 0 | 0 | 0 | 0 | 41 | 29 | 6 | 2 | 4 |
| **mF84** | 8 | 126 | 13 | 6 | 1 | 1 | 5 | 113 | 70 | 24 | 15 | 4 |
| **mF93** | 2 | 170 | 18 | 9 | 1 | 0 | 8 | 152 | 87 | 55 | 7 | 3 |
| **mK01** | 3 | 18 | 1 | 1 | 0 | 0 | 0 | 17 | 13 | 0 | 4 | 0 |
| **mO251** | 2 | 6 | 0 | 0 | 0 | 0 | 0 | 6 | 2 | 1 | 2 | 1 |
| **mP101** | 1 | 8 | 0 | 0 | 0 | 0 | 0 | 8 | 1 | 5 | 2 | 0 |
| **mP20** | 4 | 165 | 6 | 0 | 0 | 3 | 3 | 159 | 106 | 15 | 21 | 17 |
| **mP23** | 3 | 79 | 2 | 0 | 0 | 0 | 2 | 77 | 44 | 16 | 13 | 4 |
| **mV99** | 4 | 9 | 0 | 0 | 0 | 0 | 0 | 9 | 5 | 1 | 3 | 0 |
| **Total** | **78** | **3189** | **277** | **164** | **18** | **11** | **84** | **2912** | **2049** | **592** | **179** | **92** |

**Table S3.** Units recorded from CA1

|  |  |  | **MM** |  |  |  |  | **SM** |  |  |  |  |
| --- | --- | --- | --- | --- | --- | --- | --- | --- | --- | --- | --- | --- |
| **Animal ID** | **Sessions** | **Units** | **Total** | **PYR** | **INT** | **Punit** | **BIP** | **Total** | **PYR** | **INT** | **Punit** | **BIP** |
| **mA234** | 26 | 1699 | 162 | 108 | 0 | 0 | 54 | 1537 | 1276 | 191 | 35 | 35 |
| **mC41** | 32 | 2326 | 366 | 254 | 3 | 1 | 108 | 1960 | 1536 | 285 | 88 | 51 |
| **mDS1** | 17 | 803 | 63 | 58 | 0 | 0 | 5 | 740 | 645 | 74 | 18 | 3 |
| **mDS2** | 3 | 195 | 12 | 6 | 1 | 0 | 5 | 183 | 116 | 42 | 20 | 5 |
| **mK01** | 5 | 49 | 13 | 12 | 1 | 0 | 0 | 36 | 26 | 8 | 2 | 0 |
| **mO251** | 4 | 46 | 8 | 6 | 2 | 0 | 0 | 38 | 28 | 9 | 0 | 1 |
| **mP101** | 5 | 223 | 30 | 19 | 0 | 0 | 11 | 193 | 150 | 23 | 7 | 13 |
| **mP23** | 23 | 587 | 34 | 25 | 1 | 0 | 8 | 553 | 462 | 65 | 13 | 13 |
| **mV99** | 11 | 43 | 1 | 0 | 0 | 1 | 0 | 42 | 19 | 19 | 3 | 1 |
| **Total** | **126** | **5971** | **689** | **488** | **8** | **2** | **191** | **5282** | **4258** | **716** | **186** | **122** |

**Table S4.** Units recorded from CA1 on the linear track

|  |  |  |  | MM |  |  |  |  | SM |  |  |  |  |
| --- | --- | --- | --- | --- | --- | --- | --- | --- | --- | --- | --- | --- | --- |
| Animal ID | Sessions | Units | Active units<sup>1</sup> | Total | PYR | INT | Punit | BIP | Total | PYR | INT | Punit | BIP |
| **mA234** | 24 | 1633 | 572 | 20 | 6 | 0 | 0 | 14 | 552 | 402 | 124 | 15 | 11 |
| **mC41** | 32 | 2326 | 737 | 42 | 26 | 1 | 0 | 15 | 695 | 485 | 175 | 25 | 10 |
| **mDS1** | 5 | 237 | 58 | 1 | 1 | 0 | 0 | 0 | 57 | 45 | 12 | 0 | 0 |
| **mP23** | 14 | 336 | 104 | 1 | 1 | 0 | 0 | 0 | 103 | 76 | 26 | 1 | 0 |
| **Total** | **75** | **4515** | **1471** | **64** | **34** | **1** | **0** | **29** | **1407** | **1008** | **335** | **41** | **21** |

888 <sup>1</sup> Active and stable on the linear track.

## Supplementary figures

**Figure S1.**
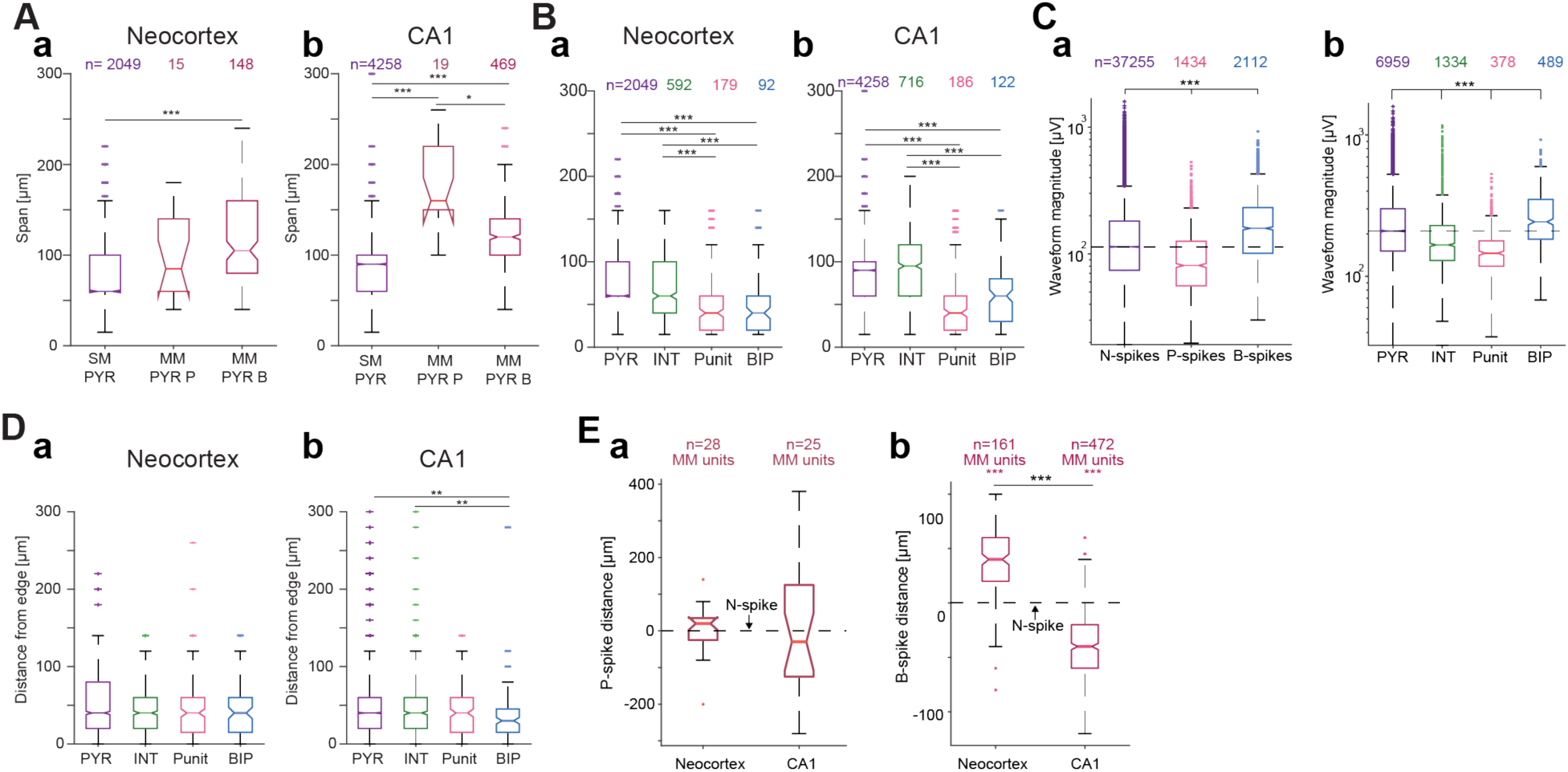
Compared with PYRs and INTs, Punits and BIPs are more spatially compact. **(A)** SM PYRs are more compact than MM PYRs in neocortex (**a**) and CA1 (**b**). Here and in **B**, **C**, and **D**, */**/***: p<0.05/p<0.01/p<0.001, Kruskal-Wallis test, corrected for multiple comparisons. **(B)** Among SM units, Punits and BIPs are more compact in space than PYRs and INTs in both neocortex (**a**) and CA1 (**b**). **(C)** (**a**) B-spikes have larger waveform magnitude compared with N-spikes, and N-spikes have larger waveform magnitude compared with P-spikes. (**b**) Waveform magnitude at main channel for every group. Magnitudes differ between all groups. Specifically, BIP waveform magnitude is larger when compared with PYRs and INTs, whereas Punit waveform magnitude is smaller compared with PYRs and INTs. **(D)** Distance between main channel and closest edge of the recording shank for SM units. Compared to PYRs and INTs, the main channel of SM BIPs in CA1 is more frequently positioned closer to the shank edge in CA1. **(E)** (**a**) In neocortex, P-spikes appear above the same-unit N-spike (median [IQR] distance: 20 [-30 30] μm). In CA1, the P-spikes appear below the N-spikes, with a distance of -20 [-140 120] μm. (**b**) B-spikes appear above the same-unit N-spike in neocortex (40 [20 60] μm). In CA1, B-spikes appear below the same-unit N-spike (−40 [-60 -20] μm). ***: p<0.001, Wilcoxon’s test. Lined ***: p<0.001, U-test.

**Figure S2.**
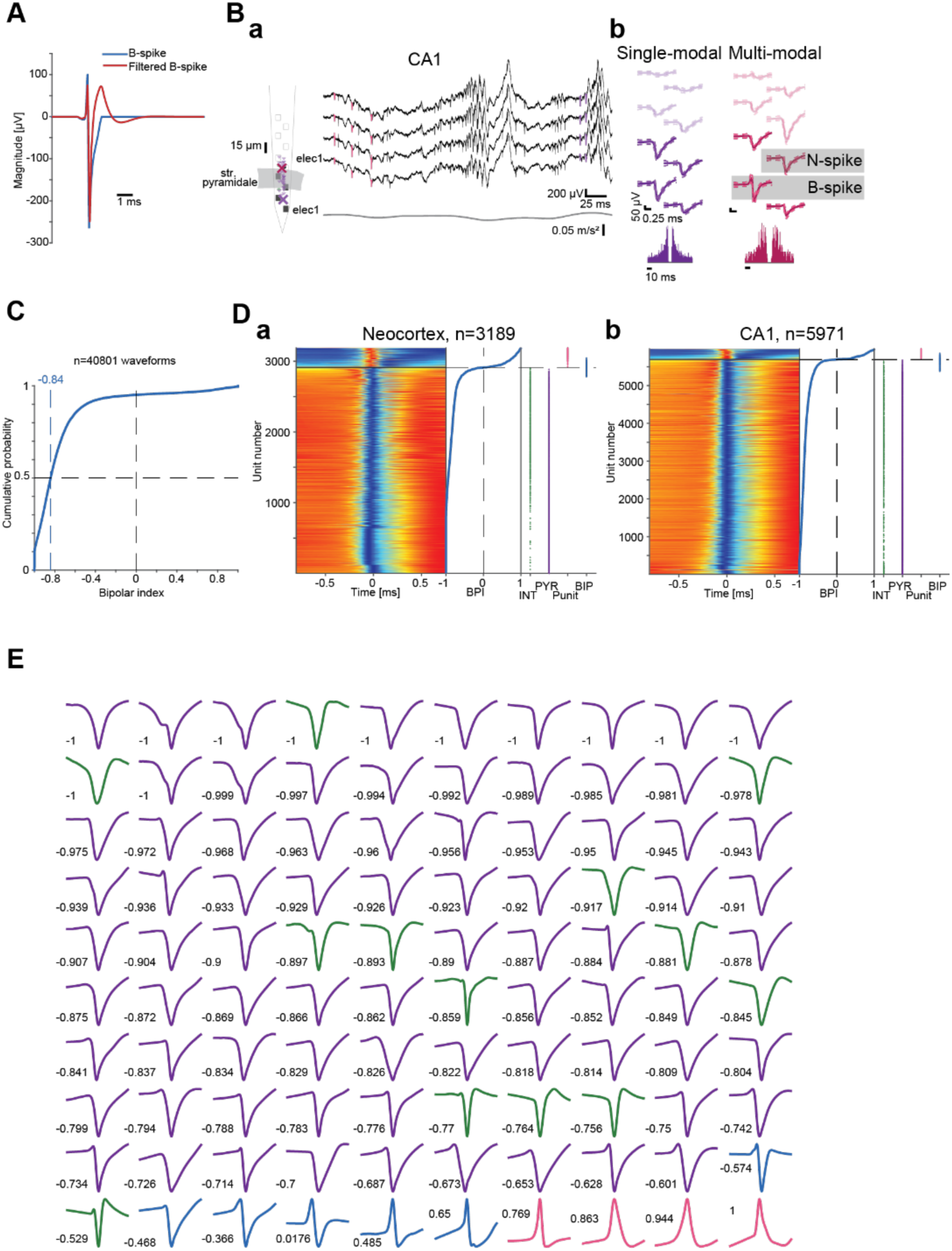
Spike waveform categorization. **(A)** High-pass filtering transforms a wideband B-spike into a triphasic spike. The specific filter used is a third-order Butterworth, 300-6000 Hz. Triphasic spikes are generated with a wide range of filter settings (e.g., high-pass at 300, 500, 800 Hz; 1^st^, 2^nd^, 3^rd^ order). **(B)** A SM PYR and a MM unit with biphasic spikes (B-spikes) recorded simultaneously from CA1 of a freely-moving mouse. (**a**) **Left**, Schematic shank with 32 simultaneously-recorded units. (**b**) Wideband waveforms and ACHs. All conventions are the same as in **Fig. 1A**. **(C)** Bipolar index (BPI) of the main channel waveforms of all 9160 units in the dataset. Blue dashed line indicates the median BPI. Most waveforms are N-spikes. **(D)** Main channel waveforms. (**a**) **Left**, Waveforms of all neocortical units. **Middle**, BPI of every unit. **Right**, Classification of each unit. (**b**) Same, for CA1 units. **(E)** Waveform examples. A statistically representative set of 100 waveforms was randomly selected from the 9160 units. Numbers indicate BPI, and colors indicate the class of the unit: PYR, purple; INT, green; BIP, blue; Punit, pink.

**Figure S3.**
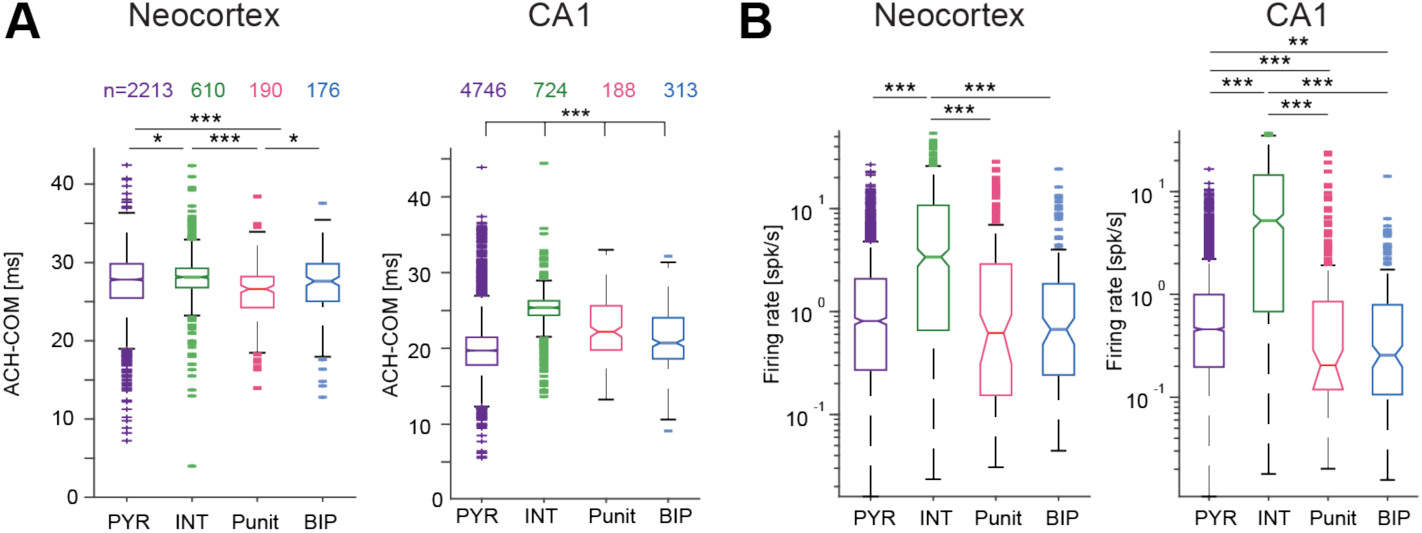
CA1 Punits and BIPs are less bursty than PYRs but more bursty than INTs. **(A)** In the neocortex, Punits are more bursty than INTs and PYRs. In CA1, PYRs are most bursty, followed by BIPs, Punits, and INTs. Sample size is indicated on top and is the same here and in **B**. Here and in **B**, ***: p<0.001, Kruskal-Wallis test, corrected for multiple comparisons. **(B)** In both neocortex and CA1, the firing rates of Punits and BIPs are lower than INTs.

**Figure S4.**
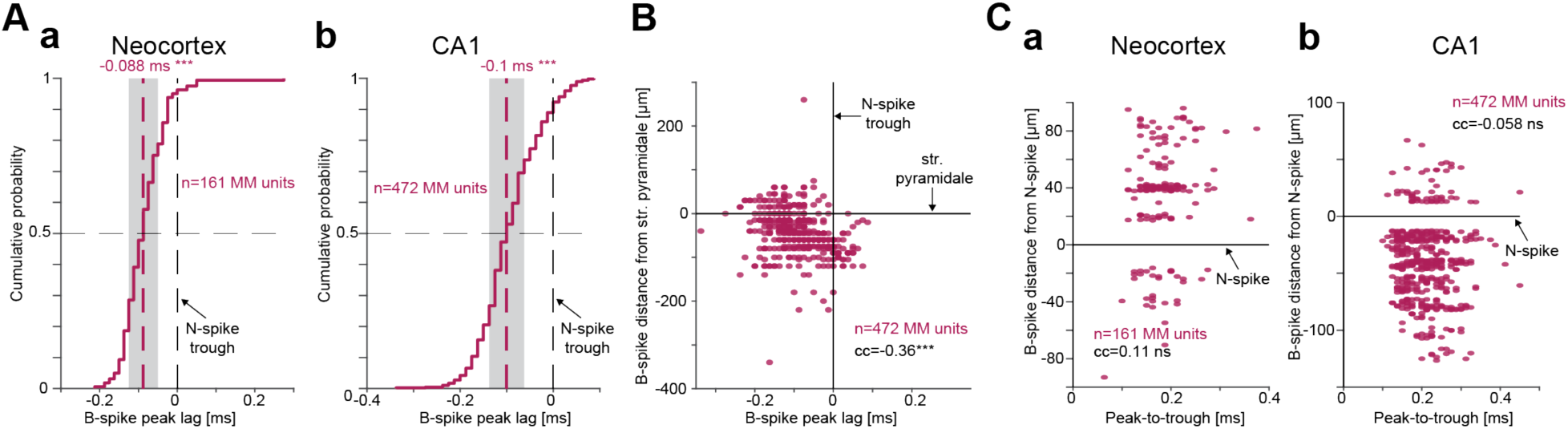
B-spike peak precedes same-unit N-spike trough. **(A)** B-spike peak precedes same-unit N-spike trough. (**a**) Time lag between B- and N-spikes in neocortical MM units. Here and in **b**: ***, p<0.001, Wilcoxon’s test comparing to a zero null. Grey patch, 95% confidence limits. (**b**) Time lag between B- and N-spike in CA1 MM units. **(B)** Distance of B-spike from the center pf CA1 str. pyramidale vs. B-spike peak lag from N-spike trough. Distances are positive when the B-spike is closer to the surface of the brain. cc, rank correlation coefficient; ***: p<0.001, permutation test. **(C)** Distance of B-spike from and N-spike vs. B-spike peak-to-trough duration in neocortex (**a**) and in CA1 (**b**).

**Figure S5.**
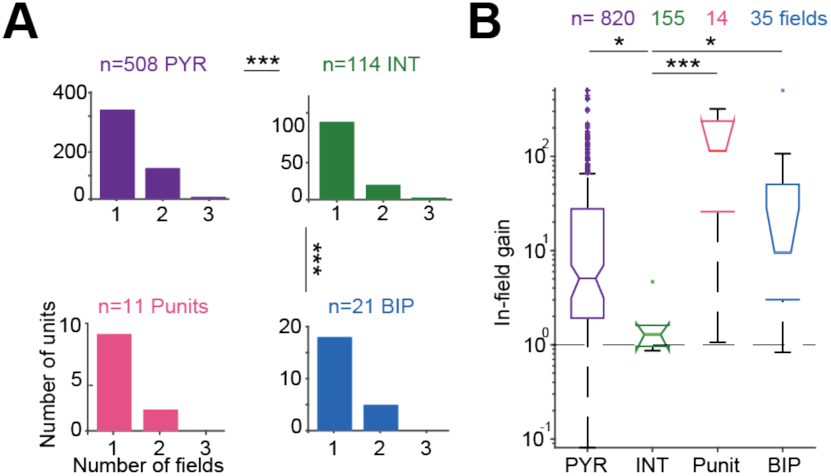
BIP gain and number of fields are not consistently different from PYR. **(A)** Number of place fields per unit. ***: p<0.001, G-test, corrected for multiple comparisons. **(B)** In-field gain. */***: p<0.05/p<0.001, Kruskal-Wallis test.

## References

1. Adrian, E.D., and Moruzzi, G. (1939). Impulses in the pyramidal tract. J. Physiol. 97, 153–199. 10.1113/jphysiol.1939.sp003798.

2. Steinmetz, N.A., Aydin, C., Lebedeva, A., Okun, M., Pachitariu, M., Bauza, M., Beau, M., Bhagat, J., Böhm, C., Broux, M., et al. (2021). Neuropixels 2.0: A miniaturized high-density probe for stable, long-term brain recordings. Science 372, eabf4588. 10.1126/science.abf4588.

3. Tranchina, D., and Nicholson, C. (1986). A model for the polarization of neurons by extrinsically applied electric fields. Biophys. J. 50, 1139–1156. 10.1016/S0006-3495(86)83558-5.

4. Buzsáki, G., Anastassiou, C.A., and Koch, C. (2012). The origin of extracellular fields and currents — EEG, ECoG, LFP and spikes. Nat. Rev. Neurosci. 13, 407–420. 10.1038/nrn3241.

5. Einevoll, G.T., Kayser, C., Logothetis, N.K., and Panzeri, S. (2013). Modelling and analysis of local field potentials for studying the function of cortical circuits. Nat. Rev. Neurosci. 14, 770–785. 10.1038/nrn3599.

6. Schomburg, E.W., Anastassiou, C.A., Buzsaki, G., and Koch, C. (2012). The spiking component of oscillatory extracellular potentials in the rat hippocampus. J. Neurosci. 32, 11798–11811. 10.1523/JNEUROSCI.0656-12.2012.

7. Deligkaris, K., Bullmann, T., and Frey, U. (2016). Extracellularly recorded somatic and neuritic signal shapes and classification algorithms for high-density microelectrode array electrophysiology. Front. Neurosci. 10, 421. 10.3389/fnins.2016.00421.

8. Gold, C., Henze, D.A., Koch, C., and Buzsáki, G. (2006). On the origin of the extracellular action potential waveform: a modeling study. J. Neurophysiol. 95, 3113–3128. 10.1152/jn.00979.2005.

9. Holt, G.R., and Koch, C. (1999). Electrical interactions via the extracellular potential near cell bodies. J. Comput. Neurosci. 6, 169–184.

10. Rall, W., and Shepherd, G.M. (1968). Theoretical reconstruction of field potentials and dendrodendritic synaptic interactions in olfactory bulb. J. Neurophysiol. 31, 884–915.

11. Matthews, R.T., and Lee, W.L. (1991). A comparison of extracellular and intracellular recordings from medial septum/diagonal band neurons in vitro. Neuroscience 42, 451–462. 10.1016/0306-4522(91)90388-5.

12. Henze, D.A., Borhegyi, Z., Csicsvari, J., Mamiya, A., Harris, K.D., and Buzsáki, G. (2000). Intracellular features predicted by extracellular recordings in the hippocampus in vivo. J. Neurophysiol. 84, 390–400. 10.1152/jn.2000.84.1.390.

13. O’Keefe, J., and Recce, M.L. (1993). Phase relationship between hippocampal place units and the EEG theta rhythm. Hippocampus 3, 317–330. 10.1002/hipo.450030307.

14. Bordi, F., and LeDoux, J.E. (1994). Response properties of single units in areas of rat auditory thalamus that project to the amygdala. Exp. Brain Res. 98, 275–286. 10.1007/BF00228415.

15. Bestel, R., van Rienen, U., Thielemann, C., and Appali, R. (2021). Influence of neuronal morphology on the shape of extracellular recordings with microelectrode arrays: a finite element analysis. IEEE Trans. Biomed. Eng. 68, 1317–1329. 10.1109/TBME.2020.3026635.

16. Rall, W. (1962). Electrophysiology of a dendritic neuron model. Biophys. J. 2, 145–167. 10.1016/S0006-3495(62)86953-7.

17. Rasminsky, M. (1980). Ephaptic transmission between single nerve fibres in the spinal nerve roots of dystrophic mice. J. Physiol. 305, 151–169.

18. Miyakawa, H., and Kato, H. (1986). Active properties of dendritic membrane examined by current source density analysis in hippocampal CA1 pyramidal neurons. Brain Res. 399, 303–309. 10.1016/0006-8993(86)91520-9.

19. Bakkum, D.J., Obien, M.E.J., Radivojevic, M., Jäckel, D., Frey, U., Takahashi, H., and Hierlemann, A. (2019). The axon initial segment is the dominant contributor to the neuron’s extracellular electrical potential landscape. Adv. Biosyst. 3, 1800308. 10.1002/adbi.201800308.

20. Emmenegger, V., Obien, M.E.J., Franke, F., and Hierlemann, A. (2019). Technologies to study action potential propagation with a focus on HD-MEAs. Front. Cell. Neurosci. 13, 159.

21. Gold, C., Girardin, C.C., Martin, K.A.C., and Koch, C. (2009). High-amplitude positive spikes recorded extracellularly in cat visual cortex. J. Neurophysiol. 102, 3340–3351. 10.1152/jn.91365.2008.

22. Barthó, P., Slézia, A., Mátyás, F., Faradzs-Zade, L., Ulbert, I., Harris, K.D., and Acsády, L. (2014). Ongoing network state controls the length of sleep spindles via inhibitory activity. Neuron 82, 1367– 1379. 10.1016/j.neuron.2014.04.046.

23. Sun, Almasi, A., Yunzab, M., Zehra, S., Hicks, D.G., Kameneva, T., Ibbotson, M.R., and Meffin, H. (2021). Analysis of extracellular spike waveforms and associated receptive fields of neurons in cat primary visual cortex. J. Physiol. 599, 2211–2238. 10.1113/JP280844.

24. Tovar, K.R., Bridges, D.C., Wu, B., Randall, C., Audouard, M., Jang, J., Hansma, P.K., and Kosik, K.S. (2018). Action potential propagation recorded from single axonal arbors using multielectrode arrays. J. Neurophysiol. 120, 306–320. 10.1152/jn.00659.2017.

25. Lewandowska, M.K., Bakkum, D.J., Rompani, S.B., and Hierlemann, A. (2015). Recording large extracellular spikes in microchannels along many axonal sites from individual neurons. PLoS ONE 10, e0118514. 10.1371/journal.pone.0118514.

26. Shein-Idelson, M., Pammer, L., Hemberger, M., and Laurent, G. (2017). Large-scale mapping of cortical synaptic projections with extracellular electrode arrays. Nat. Methods 14, 882–890. 10.1038/nmeth.4393.

27. Raastad, M., and Shepherd, G.M.G. (2003). Single-axon action potentials in the rat hippocampal cortex. J. Physiol. 548, 745–752. 10.1113/jphysiol.2002.032706.

28. Robbins, A.A., Fox, S.E., Holmes, G.L., Scott, R.C., and Barry, J.M. (2013). Short duration waveforms recorded extracellularly from freely moving rats are representative of axonal activity. Front. Neural Circuits 7, 181. 10.3389/fncir.2013.00181.

29. Sibille, J., Gehr, C., Benichov, J.I., Balasubramanian, H., Teh, K.L., Lupashina, T., Vallentin, D., and Kremkow, J. (2022). High-density electrode recordings reveal strong and specific connections between retinal ganglion cells and midbrain neurons. Nat. Commun. 13, 5218. 10.1038/s41467-022-32775-2.

30. Zhu, S., Xia, R., Chen, X., and Moore, T. (2020). Heterogeneity of neuronal populations within columns of primate V1 revealed by high-density recordings (Neuroscience) 10.1101/2020.12.22.424048.

31. Paulk, A.C., Kfir, Y., Khanna, A.R., Mustroph, M.L., Trautmann, E.M., Soper, D.J., Stavisky, S.D., Welkenhuysen, M., Dutta, B., Shenoy, K.V., Hochberg, L.R., Richardson, R.M., Williams, Z.M., Cash S.S. (2022). Large-scale neural recordings with single neuron resolution using Neuropixels probes in human cortex. Nat. Neurosci. 25, 252–263. 10.1038/s41593-021-00997-0.

32. Claverol-Tinture, E., and Pine, J. (2002). Extracellular potentials in low-density dissociated neuronal cultures. J. Neurosci. Methods 117, 13–21. 10.1016/S0165-0270(02)00043-2.

33. Gold, C., Henze, D.A., and Koch, C. (2007). Using extracellular action potential recordings to constrain compartmental models. J. Comput. Neurosci. 23, 39–58. 10.1007/s10827-006-0018-2.

34. Bereshpolova, Y., Amitai, Y., Gusev, A.G., Stoelzel, C.R., and Swadlow, H.A. (2007). Dendritic backpropagation and the state of the awake neocortex. J. Neurosci. 27, 9392–9399. 10.1523/JNEUROSCI.2218-07.2007.

35. Buzsaki, G., Penttonen, M., Nadasdy, Z., and Bragin, A. (1996). Pattern and inhibition-dependent invasion of pyramidal cell dendrites by fast spikes in the hippocampus in vivo. Proc. Natl. Acad. Sci. 93, 9921–9925. 10.1073/pnas.93.18.9921.

36. Jia, X., Siegle, J.H., Bennett, C., Gale, S.D., Denman, D.J., Koch, C., and Olsen, S.R. (2019). High-density extracellular probes reveal dendritic backpropagation and facilitate neuron classification. J. Neurophysiol. 121, 1831–1847. 10.1152/jn.00680.2018.

37. Stuart, Spruston, N., Sakmann, B., and Häusser, M. (1997). Action potential initiation and backpropagation in neurons of the mammalian CNS. Trends Neurosci. 20, 125–131. 10.1016/S0166-2236(96)10075-8.

38. Golding, N.L., Kath, W.L., and Spruston, N. (2001). Dichotomy of action-potential backpropagation in CA1 pyramidal neuron dendrites. J. Neurophysiol. 86, 2998–3010. 10.1152/jn.2001.86.6.2998.

39. Kole, M.H.P., Letzkus, J.J., and Stuart, G.J. (2007). Axon initial segment Kv1 channels control axonal action potential waveform and synaptic efficacy. Neuron 55, 633–647. 10.1016/j.neuron.2007.07.031.

40. Migliore, M., and Shepherd, G.M. (2002). Emerging rules for the distributions of active dendritic conductances. Nat. Rev. Neurosci. 3, 362–370. 10.1038/nrn810.

41. Stuart, and Sakmann, B. (1994). Active propagation of somatic action potentials into neocortical pyramidal cell dendrites. Nature 367, 69–72. 10.1038/367069a0.

42. Szabadics, J., Varga, C., Molnár, G., Oláh, S., Barzó, P., and Tamás, G. (2006). Excitatory effect of GABAergic axo-axonic cells in cortical microcircuits. Science 311, 233–235. 10.1126/science.1121325.

43. Koos, T., Tecuapetla, F., and Tepper, J.M. (2011). Glutamatergic signaling by midbrain dopaminergic neurons: recent insights from optogenetic, molecular and behavioral studies. Curr. Opin. Neurobiol. 21, 393–401. 10.1016/j.conb.2011.05.010.

44. Trudeau, L.-E., Hnasko, T.S., Wallén-Mackenzie, Å., Morales, M., Rayport, S., and Sulzer, D. (2014). Chapter 6 - The multilingual nature of dopamine neurons. In Progress in Brain Research Dopamine., M. Diana, G. Di Chiara, and P. Spano, eds. (Elsevier), pp. 141–164. 10.1016/B978-0-444-63425-2.00006-4.

45. O’Donohue, T.L., Millington, W.R., Handelmann, G.E., Contreras, P.C., and Chronwall, B.M. (1985). On the 50th anniversary of Dale’s law: multiple neurotransmitter neurons. Trends Pharmacol. Sci. 6, 305–308. 10.1016/0165-6147(85)90141-5.

46. Dale, H. (1935). Pharmacology and nerve-endings. Proc. R. Soc. Med. 28, 319–332. 10.1177/003591573502800330.

47. Levi, A., Spivak, L., Sloin, H.E., Someck, S., and Stark, E. (2022). Error correction and improved precision of spike timing in converging cortical networks. Cell Rep. 40, 111383. 10.1016/j.celrep.2022.111383.

48. Skaggs, W.E., McNaughton, B.L., Wilson, M.A., and Barnes, C.A. (1996). Theta phase precession in hippocampal neuronal populations and the compression of temporal sequences. Hippocampus 6, 149–172. 10.1002/(SICI)1098-1063(1996)6:2<149::AID-HIPO6>3.0.CO;2-K.

49. Stark, E., Roux, L., Eichler, R., Senzai, Y., Royer, S., and Buzsáki, G. (2014). Pyramidal cell-interneuron interactions underlie hippocampal ripple oscillations. Neuron 83, 467–480. 10.1016/j.neuron.2014.06.023.

50. Destexhe, A., and Bedard, C. (2012). Do neurons generate monopolar current sources? J. Neurophysiol. 108, 953–955. 10.1152/jn.00357.2012.

51. Golding, N.L., and Spruston, N. (1998). Dendritic sodium spikes are variable triggers of axonal action potentials in hippocampal CA1 pyramidal neurons. Neuron 21, 1189–1200. 10.1016/S0896-6273(00)80635-2.

52. Sun, Q., Srinivas, K.V., Sotayo, A., and Siegelbaum, S.A. (2014). Dendritic Na+ spikes enable cortical input to drive action potential output from hippocampal CA2 pyramidal neurons. eLife 3, e04551. 10.7554/eLife.04551.

53. Losonczy, A., and Magee, J.C. (2006). Integrative properties of radial oblique dendrites in hippocampal CA1 pyramidal neurons. Neuron 50, 291–307. 10.1016/j.neuron.2006.03.016.

54. Schiller, J., Schiller, Y., Stuart, G., and Sakmann, B. (1997). Calcium action potentials restricted to distal apical dendrites of rat neocortical pyramidal neurons. J. Physiol. 505, 605–616. 10.1111/j.1469-7793.1997.605ba.x.

55. Gulledge, A.T., Kampa, B.M., and Stuart, G.J. (2005). Synaptic integration in dendritic trees. J. Neurobiol. 64, 75–90. 10.1002/neu.20144.

56. Tomassy, G.S., Berger, D.R., Chen, H.-H., Kasthuri, N., Hayworth, K.J., Vercelli, A., Seung, H.S., Lichtman, J.W., and Arlotta, P. (2014). Distinct profiles of myelin distribution along single axons of pyramidal neurons in the neocortex. Science 344, 319–324. 10.1126/science.1249766.

57. Stadelmann, C., Timmler, S., Barrantes-Freer, A., and Simons, M. (2019). Myelin in the central nervous system: structure, function, and pathology. Physiol. Rev. 99, 1381–1431. 10.1152/physrev.00031.2018.

58. Micheva, K.D., Wolman, D., Mensh, B.D., Pax, E., Buchanan, J., Smith, S.J., and Bock, D.D. (2016). A large fraction of neocortical myelin ensheathes axons of local inhibitory neurons. eLife 5, e15784. 10.7554/eLife.15784.

59. Lorincz, A., and Nusser, Z. (2008). Cell-type-dependent molecular composition of the axon initial segment. J. Neurosci. 28, 14329–14340. 10.1523/JNEUROSCI.4833-08.2008.

60. Dugué, G.P., Brunel, N., Hakim, V., Schwartz, E., Chat, M., Lévesque, M., Courtemanche, R., Léna, C., and Dieudonné, S. (2009). Electrical coupling mediates tunable low-frequency oscillations and resonance in the cerebellar golgi cell network. Neuron 61, 126–139. 10.1016/j.neuron.2008.11.028.

61. van Welie, I., Roth, A., Ho, S.S.N., Komai, S., and Häusser, M. (2016). Conditional spike transmission mediated by electrical coupling ensures millisecond precision-correlated activity among interneurons in vivo. Neuron 90, 810–823. 10.1016/j.neuron.2016.04.013.

62. Gibson, J.R., Beierlein, M., and Connors, B.W. (1999). Two networks of electrically coupled inhibitory neurons in neocortex. Nature 402, 75–79. 10.1038/47035.

63. Curti, S., Davoine, F., and Dapino, A. (2022). Function and plasticity of electrical synapses in the mammalian brain: role of non-junctional mechanisms. Biology 11, 81. 10.3390/biology11010081.

64. Cid, E., and de la Prida, L.M. (2019). Methods for single-cell recording and labeling in vivo. J. Neurosci. Methods 325, 108354. 10.1016/j.jneumeth.2019.108354.

65. Gilman, J.P., Medalla, M., and Luebke, J.I. (2017). Area-specific features of pyramidal neurons—a comparative study in mouse and rhesus monkey. Cereb. Cortex 27, 2078–2094. 10.1093/cercor/bhw062.

66. Reuben, J.P., Werman, R., and Grundfest, H. (1961). The ionic mechanisms of hyperpolarizing responses in lobster muscle fibers. J. Gen. Physiol. 45, 243–265. 10.1085/jgp.45.2.243.

67. del Castillo, J., and Morales, T. (1967). Extracellular action potentials recorded from the interior of the giant esophageal cell of ascaris. J. Gen. Physiol. 50, 631–645. 10.1085/jgp.50.3.631.

68. Izhikevich, E.M. (2007). Dynamical systems in neuroscience (MIT Press).

69. de Cheveigné, A., and Nelken, I. (2019). Filters: when, why, and how (not) to use them. Neuron 102, 280–293. 10.1016/j.neuron.2019.02.039.

70. Quiroga, Q.R. (2009). What is the real shape of extracellular spikes? J. Neurosci. Methods 177, 194–198. 10.1016/j.jneumeth.2008.09.033.

71. Mizuseki, K., Royer, S., Diba, K., and Buzsáki, G. (2012). Activity dynamics and behavioral correlates of CA3 and CA1 hippocampal pyramidal neurons. Hippocampus 22, 1659–1680. 10.1002/hipo.22002.

72. Schmidt, H., Gour, A., Straehle, J., Boergens, K.M., Brecht, M., and Helmstaedter, M. (2017). Axonal synapse sorting in medial entorhinal cortex. Nature 549, 469–475. 10.1038/nature24005.

73. Stuart, G., Spruston, N., and Häusser, M. (2007). Dendrites (Oxford University Press).

74. Hierlemann, A., Frey, U., Hafizovic, S., and Heer, F. (2011). Growing cells atop microelectronic chips: interfacing electrogenic cells in vitro with CMOS-based microelectrode arrays. Proc. IEEE 99, 252–284. 10.1109/JPROC.2010.2066532.

75. Boiko, T., Rasband, M.N., Levinson, S.R., Caldwell, J.H., Mandel, G., Trimmer, J.S., and Matthews, G. (2001). Compact myelin dictates the differential targeting of two sodium channel isoforms in the same axon. Neuron 30, 91–104. 10.1016/S0896-6273(01)00265-3.

76. Rosker, C., Lohberger, B., Hofer, D., Steinecker, B., Quasthoff, S., and Schreibmayer, W. (2007). The TTX metabolite 4,9-anhydro-TTX is a highly specific blocker of the Nav1.6 voltage-dependent sodium channel. Am. J. Physiol.-Cell Physiol. 293, C783–C789. 10.1152/ajpcell.00070.2007.

77. Leterrier, C., Brachet, A., Fache, M.-P., and Dargent, B. (2010). Voltage-gated sodium channel organization in neurons: Protein interactions and trafficking pathways. Neurosci. Lett. 486, 92–100. 10.1016/j.neulet.2010.08.079.

78. Waxman, S., and Bennett, M. (1972). Relative conduction velocities of small myelinated and non-myelinated fibres in the central nervous system. Nature. New Biol. 238, 217–219. 10.1038/newbio238217a0.

79. Telfeian, A.E., and Connors, B.W. (2003). Widely integrative properties of layer 5 pyramidal cells support a role for processing of extralaminar synaptic inputs in rat neocortex. Neurosci. Lett. 343, 121–124. 10.1016/S0304-3940(03)00379-3.

80. Hirsch, J., and Gilbert, C. (1991). Synaptic physiology of horizontal connections in the cat’s visual cortex. J. Neurosci. 11, 1800–1809. 10.1523/JNEUROSCI.11-06-01800.1991.

81. Micheva, K.D., Kiraly, M., Perez, M.M., and Madison, D.V. (2021). Conduction velocity along the local axons of parvalbumin interneurons correlates with the degree of axonal myelination. Cereb. Cortex 31, 3374–3392. 10.1093/cercor/bhab018.

82. Benucci, A., Frazor, R.A., and Carandini, M. (2007). Standing waves and traveling waves distinguish two circuits in visual cortex. Neuron 55, 103–117. 10.1016/j.neuron.2007.06.017.

83. Vickers, J.C., King, A.E., Woodhouse, A., Kirkcaldie, M.T., Staal, J.A., McCormack, G.H., Blizzard, C.A., Musgrove, R.E.J., Mitew, S., Liu, Y., Chuckowree J.A., Bibari O., Dickson T.C. (2009). Axonopathy and cytoskeletal disruption in degenerative diseases of the central nervous system. Brain Res. Bull. 80, 217–223. 10.1016/j.brainresbull.2009.08.004.

84. Ferguson, B. (1997). Axonal damage in acute multiple sclerosis lesions. Brain 120, 393–399. 10.1093/brain/120.3.393.

85. Brown, A., McFarlin, D.E., and Raine, C.S. (1982). Chronologic neuropathology of relapsing experimental allergic encephalomyelitis in the mouse. Lab. Investig. J. Tech. Methods Pathol. 46, 171– 185.

86. Sloin, H.E., Bikovski, L., Levi, A., Amber-Vitos, O., Katz, T., Spivak, L., Someck, S., Gattegno, R., Sivroni, S., Sjulson, L., Stark E. (2022). Hybrid offspring of C57BL/6J mice exhibit improved properties for neurobehavioral research. eNeuro 9. 10.1523/ENEURO.0221-22.2022.

87. Stark, E., Koos, T., and Buzsáki, G. (2012). Diode probes for spatiotemporal optical control of multiple neurons in freely moving animals. J. Neurophysiol. 108, 349–363. 10.1152/jn.00153.2012.

88. Noked, O., Levi, A., Someck, S., Amber-Vitos, O., and Stark, E. (2021). Bidirectional optogenetic control of inhibitory neurons in freely-moving mice. IEEE Trans. Biomed. Eng. 68, 416–427. 10.1109/TBME.2020.3001242.

89. Gaspar, N., Eichler, R., and Stark, E. (2019). A novel low-noise movement tracking system with real-time analog output for closed-loop experiments. J. Neurosci. Methods 318, 69–77. 10.1016/j.jneumeth.2018.12.016.

90. Rossant, C., Kadir, S.N., Goodman, D.F.M., Schulman, J., Hunter, M.L.D., Saleem, A.B., Grosmark, A., Belluscio, M., Denfield, G.H., Ecker, A.S., Tolias A.S., Solomon S., Buzsáki G., Carandini M., Harris K.D. (2016). Spike sorting for large, dense electrode arrays. Nat. Neurosci. 19, 634–641. 10.1038/nn.4268.

91. Pachitariu, M., Steinmetz, N., Kadir, S., Carandini, M., and Harris, K.D. (2016). Kilosort: realtime spike-sorting for extracellular electrophysiology with hundreds of channels. 061481. 10.1101/061481.

92. Stark, E., Eichler, R., Roux, L., Fujisawa, S., Rotstein, H.G., and Buzsáki, G. (2013). Inhibition-induced theta resonance in cortical circuits. Neuron 80, 1263–1276. 10.1016/j.neuron.2013.09.033.

93. Spivak, L., Levi, A., Sloin, H.E., Someck, S., and Stark, E. (2022). Deconvolution improves the detection and quantification of spike transmission gain from spike trains. Commun. Biol. 5, 520. 10.1038/s42003-022-03450-5.

94. Sloin, H.E., Levi, A., Someck, S., Spivak, L., and Stark, E. (2022). High fidelity theta phase rolling of CA1 neurons. J. Neurosci. 42, 3184–3196. 10.1523/JNEUROSCI.2151-21.2022.

